# Cell division history encodes directional information of fate transitions

**DOI:** 10.1101/2022.10.06.511094

**Authors:** Kun Wang, Liangzhen Hou, Zhaolian Lu, Xin Wang, Zhike Zi, Weiwei Zhai, Xionglei He, Christina Curtis, Da Zhou, Zheng Hu

**Author notes:** Correspondence to (D.Z.); (Z.H.).

## Abstract

Single-cell RNA-sequencing (scRNA-seq) enables systematic mapping of cellular differentiation trajectories. However, inferring the cell-fate transitions under diseases or perturbations is still challenging due to the high cellular plasticity. Here, we demonstrate that monotonically expressed genes (MEGs) along cell divisions record the directions of state transitions regardless of the cellular processes. We developed a computational framework (PhyloVelo) to identify MEGs and reconstruct a novel transcriptomic velocity field by leveraging both scRNA-seq and phylogenetic information. PhyloVelo accurately recovered linear, bifurcated and convergent differentiations in simulations and *C. elegans.* It outperformed current approaches for delineating cellular trajectories in embryo development and tumor evolution through analysis of five CRISPR/Cas9-based lineage tracing datasets. Together, our study unveils an internal cellular clock and provides a powerful method for cellfate analysis in diverse biological contexts.

## Introduction

Organism development and disease progression both involve serial cell-fate transitions upon repeated cell divisions. Essentially, all cells in an organism are related by a phylogenetic tree where the root represents the zygote or disease-causing cells, the branches represent cell divisions, and the leaves represent the terminal cells at various phenotypic states (e.g. cell types) (*1–4*). To understand how cell fate is determined, it is important to identify the order of cell-state transitions acting in the lineage tree and the underlying genetic or epigenetic mechanisms that precipitate these transitions (*5*).

Single-cell RNA sequencing (scRNA-seq) has been a powerful approach to study cellular differentiating trajectories (*6–10*). However, the transcriptomic trajectories may or may not be equivalent to the true lineage paths of a progenitor population (*11–14*). One example is convergent differentiation where distinct progenitors can converge on the same terminal state (*15–19*). In this case, similar cellular states do not reflect a closer lineage relationship (*13*). Moreover, predicting the fate directions often requires prior knowledge of the initial/terminal cell types (*9*) or relies on the information of gene expression diversity during normal development (*10*). This limits their applications to normally differentiating systems (*20*), while abnormal development and disease progression often involve noncanonical cell-fate transitions such as dedifferentiation and transdifferentiation (*21*). RNA velocity (*22, 23*) provides a novel and powerful framework to predict cellular state transitions by leveraging the internal dynamics of spliced/unspliced RNAs, thus can be readily applied to the disease or perturbated conditions. However, the intrinsic high dynamics of RNA transcription and degradation often violates the constant rate assumptions in the model, which can lead to uncertain estimations (*23–25*). Taken together, cell-state transitions are challenging to distinguish using transcriptomic data alone (*11–14*).

The recent breakthrough of using CRISPR/Cas9 editing to record cell lineages offers an unprecedented opportunity to map the cell lineage tree at whole-organism or whole-organ level (*26–29*). Importantly, simultaneous analysis of single-cell transcriptomes and lineage tree makes it possible to uncover the complex developmental dynamics, as well as the molecular mechanisms, of cell fate commitment (*13, 30–35*). For instance, simultaneous single-cell lineage recording and transcriptomic profiling revealed the transcriptional convergence of endodermal cells from both extra-embryonic and embryonic origins during mouse embryogenesis (*32*). Although the significance of CRISPR lineage tracing coupled with single-cell transcriptomics has been widely acknowledged in developmental biology and somatic evolution (*13, 29, 33*), a computational framework to integrate dual information for reconstructing cellular trajectories is still lacking. Previous efforts have been made to utilize static barcoding information (e.g. LARRY system (*36*)) for quantitative cell-fate analysis such as the CoSpar algorithm (*37*). The only algorithm in the literature that has taken advantage of lineage tree for trajectory inference is LineageOT (*38*), which however, relies on timecourse scRNA-seq data and an invariant cell lineage tree. This type of data is currently only available in *C. elegans* (*14*), thus preventing its application to more common datasets such as CRISPR-based lineage tracing data.

In this study, we described a novel framework to systematically map cell-fate transitions by utilizing both single-cell transcriptomic and lineage tree information. Our method, called PhyloVelo (**Fig. 1**), exploits the MEGs along a cell lineage tree to quantify the transcriptomic velocity fields from phylogeny-resolved single-cell RNA-seq data. We have verified the capacity and robustness of PhyloVelo to resolve complex lineage structures in both normal development (*C. elegans* and mouse embryos) and disease progression (tumor initiation and metastasis) in numerous CRISPR/Cas9-based lineage tracing datasets. We surprisingly found MEGs were strongly enriched in ribosome-mediated processes across tissues (mouse embryo, mouse tumor, cancer and normal cell lines) and organisms (mouse and human), thus exposing an internal clock-like gene expression program during cell proliferation and differentiation.

**Fig. 1.**
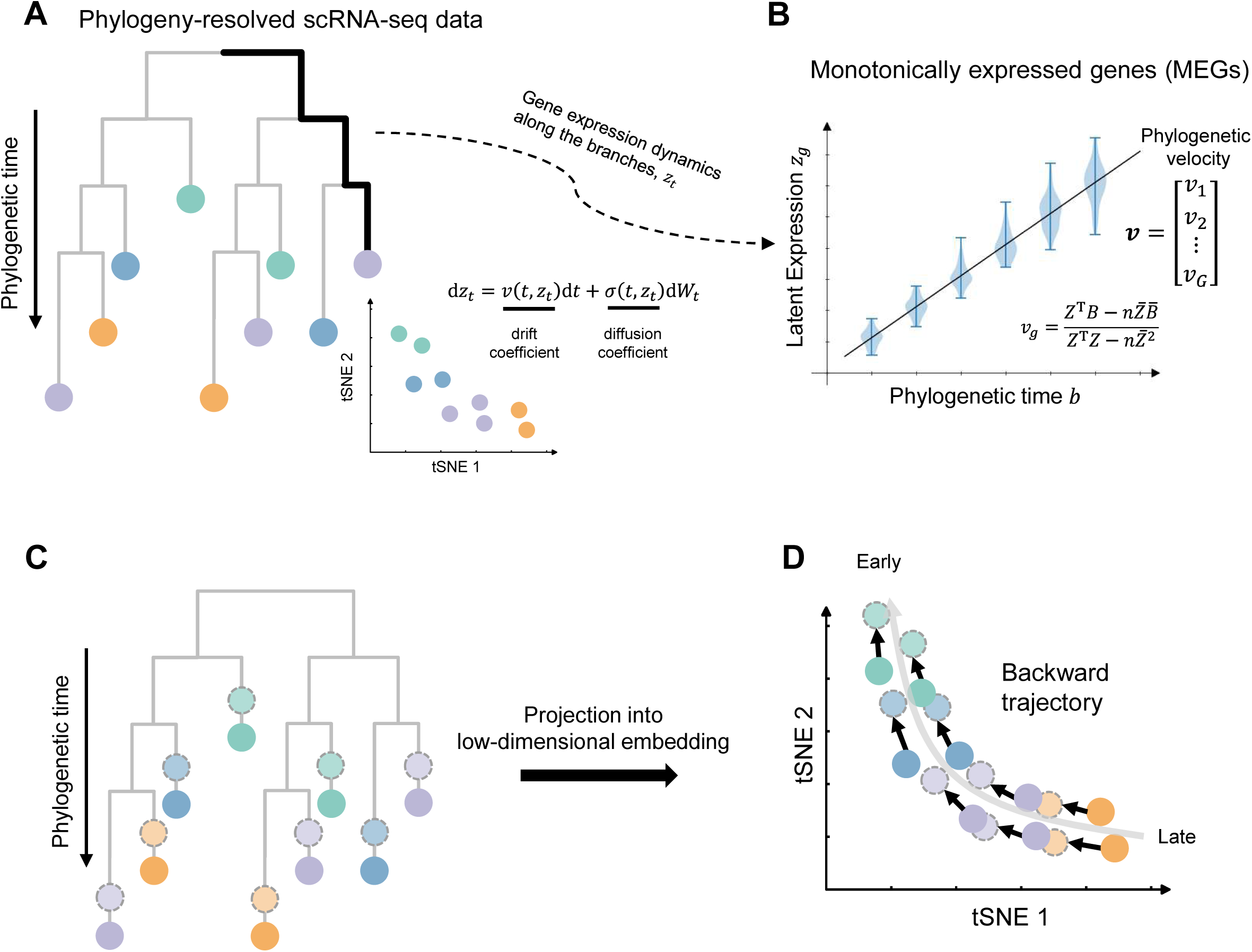
Schematic of the PhyloVelo framework. (**A**) Schematic of dual profiles of single-cell RNA-sequencing (scRNA-seq) and phylogenetic tree, namely phylogeny-resolved scRNA-seq data. The phylogenetic time corresponds to the number of cell divisions or mutations. A characteristic t-distributed Stochastic Neighbor Embedding (tSNE) of scRNA-seq data is also shown at the bottom. Colors are labeled by cell types. (**B**) An example of monotonically expressed genes (MEGs) whose latent expressions are associated with the phylogenetic distances from terminal cells to the root of the cell lineage tree. A diffusion process (also called the stochastic differential equation) is used to model the dynamics of latent gene expression *z_t_* over the phylogenetic branches. This enables the estimation of phylogenetic velocity, ***v*** = (*v*_1_, *v*_2_,⋯, *v_G_*), which corresponds to the drift coefficients of *G* MEGs in the diffusion process. (**C**) Phylogenetic velocity predicts the past transcriptional state of each cell before a unit of phylogenetic time (one cell division or mutation). (**D**) Projection of the phylogenetic velocity into low dimensional embedding enables the mapping of cell-state trajectory in a backward manner.

## Results

### A novel transcriptomic velocity field reconstructed by MEGs

RNA velocity (*22, 23*) exploits the dynamics of spliced/unspliced RNAs in single cells to estimate the time derivative (or velocity) of gene expression states (ds/dt with s representing the high-dimensional expression state and *t* representing time). This enables the reconstruction of a velocity vector field of cellular state transitions on low-dimensional embedding. We anticipated that any measurements of ds/dt can be similarly employed to establish the velocity field. For the phylogeny-resolved scRNA-seq data (**Fig. 1A**), we sought to identify MEGs whose expressions increase or decrease continuously along the phylogenetic branches regardless of the terminal cell types (**Fig. 1B**). For a MEG, the ds/dt can be estimated by the rate of its expression changes per unit of phylogenetic time (one cell division or mutation). Therefore, the state of MEGs together serves as an internal clock of transcriptomic dynamics in a proliferating cell population (**Methods**).

To identify MEGs with phylogeny-resolved scRNA-seq data (e.g. CRISPR/Cas9 lineage tracing data), we first need to estimate the latent expressions of each gene in the ancestral division history of a cell (**Fig. 1A-B**). Inspired by the classic models of continuous trait evolution in phylogenetics (*39*), we modeled the continuously varying gene expressions by a diffusion process (also called stochastic differential equation) (**Fig. 1A-B, Methods**). Each gene has a specific rate of expression changes, namely the drift coefficient *v*(*t,z_t_*) per unit of phylogenetic distance, where *t* is the phylogenetic time started from the root of tree and *z_t_* is the latent expression for a gene at time *t* (**Fig. 1B**). MEGs were identified by their significant association (Spearman’s correlation, *P*<0.05) between the latent expressions, 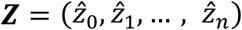, and the phylogenetic time from terminal cells to the root, ***B*** = (*b*_0_, *b*_2_,…, *b_n_*), where *n* was the cell number (**Fig. 1B**). The drift coefficients of all MEGs in a dataset, *v* = (*v*_1_, *v*_2_,⋯,*v_G_*), were thus referred to as phylogenetic velocity (or PhyloVelo) (**Fig. B**). As shown in **Fig. 1C**, phylogenetic velocity can be used to predict the past expression state of each cell before a unit of phylogenetic time *Δt* (one cell division or mutation), ***s***\* = ***s*** – ***v***Δ*t*. Similar to RNA velocity (*22, 23*), phylogenetic velocity ***v*** can also be projected into low dimensional embedding such as t-distributed Stochastic Neighbor Embedding (tSNE) or Uniform Manifold Approximation and Projection (UMAP), which generates the transcriptomic vector fields (**Fig. 1D**). Unlike RNA velocity where the velocity fields point to future extrapolated states, phylogenetic velocity fields point to the instantaneously past states (**Fig. 1D**), thus reconstructing fate-transition map in a backward manner.

### PhyloVelo robustly recovers complex lineages in simulations and *C. elegans*

We first sought to test PhyloVelo with simulation data where various lineage structures were considered, including linear, bifurcated and convergent differentiations (**Fig. 2A-C**). A lineage-imbedded scRNA-seq simulator PROSSTT (*40*) was modified to record individual cell divisions and generate single-cell UMI counts simultaneously (**Methods**). To model cell differentiations, different cell types each showing a characteristic gene expression program were simulated in the three lineage structures, respectively (**Fig 2A-C, Fig. S1**). We also simulated random mutations that occur during cell divisions, which allows to build mutation-based cell phylogeny (**Methods**). Of note, simulations showed that different cell types were highly intermixed on the lineage tree (**Fig 2A-C**), a phenomenon that appears to be general for organ development across diverse species, such as flies (*41*), zebrafish (*42*) and mice (*43*). Nevertheless, the dimensionality reduction embedding (tSNE) of simulated scRNA-seq data recapitulated the actual lineage structures (**Fig 2A-C**).

**Fig. 2.**
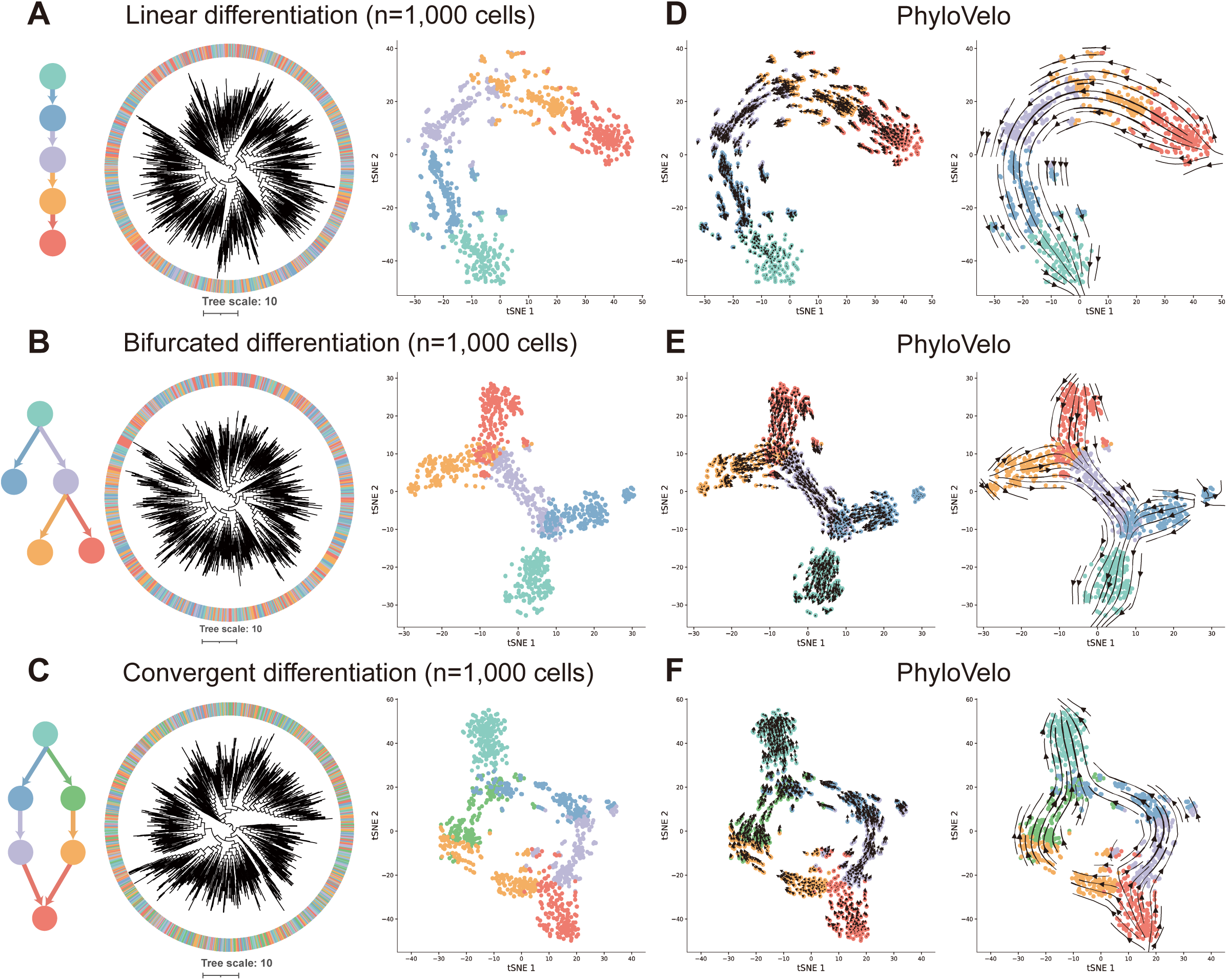
PhyloVelo recovers complex cell lineages in simulations. **Simulation of single-cell RNA-seq data and coupled cell-division history under linear** (**A**), bifurcated (**B**), and convergent (**C**) differentiation models, respectively. Colors are labeled by cell types. Each simulation consists of 1,000 cells randomly sampled from a growing cell population at 10,000 cells. Each cell has 2,000 expressed genes, including 200-300 MEGs. (**D-F**) Phylogenetic velocity fields reconstructed by PhyloVelo for the corresponding differentiation scenarios. The left panel shows the single-cell level of velocity fields, while the right panel shows the same velocity fields visualized as streamlines in scVelo.

By applying PhyloVelo to the simulation data, we first found that MEGs following either increasing or decreasing dynamics can be robustly detected with our algorithm (precision >80% for all three differentiation models) (**Fig. S2**). Second, the phylogenetic velocity fields in the tSNE diagram generate a map of stream-like state transitions in a backward manner pointing to the extrapolated past states of single cells (**Fig. 2D-F**). Third, the estimated phylogenetic velocity fields based on the mutation-based phylogenies were highly concordant with that based on the ground-truth division history (**Fig. S3-4**), although inaccurate velocity estimations in local lineages were noted when the mutation rate was rather low (mean mutation rate *u*=0.1 per cell division). Importantly, PhyloVelo inferences were robust to the phylogenetic reconstruction methods, where the directions of phylogenetic velocities using three different phylogenetic methods were highly consistent with each other and also with the ground truth (over 99% consistence for linear and bifurcated differentiations, over 75% consistence for convergent differentiation; **Fig. S5**). Fourth, while classic trajectory inference algorithms such as monocle3 (*7*), slingshot (*44*), and PAGA (*45*) can accurately identify the backbones of linear and bifurcated lineage structures, only PAGA was able to identify the circular structure in the convergent differentiation (**Fig. S6**). In fact, additional information on initial or terminal cell types is needed to define the trajectory directions using the aforementioned three algorithms. This is expected because most trajectory inference methods are inadequate for single-cell datasets containing a convergent trajectory, and also rely on prior information of initial/terminal cell types (*9*).

We next applied PhyloVelo to *C. elegans* given that the embryonic lineage tree of this organism is entirely known (*2*). The scRNA-seq data from temporal *C. elegans* embryos are also available and have been mapped to the invariant lineage tree, as described by Packer *et al.* (*14*). Thus, *C. elegans* is an ideal system to benchmark our method. We focused on the AB lineage (**Fig. 3A**), which had the densest single-cell annotations from generation 5 (32-cell stage) to 12 (threefold stage of development) (*14*), with mostly ectoderm accounting for ~70% of the terminal cells in the embryo. Since many nodes on the lineage tree have been sampled multiple times through sequencing of pooled embryos, one random cell was chosen to represent the corresponding lineage node. This resulted in 298 non-repetitive cells for the AB lineage, denoted as a single pseudo-embryo. By analyzing the correlation between latent gene expressions and cell generation times, we identified 322 significant (*P*<0.05) MEGs with 21 and 301 increasing expression and decreasing in expression, respectively (**Fig. 3B, Table S1**). This was consistent with the observed global decline in gene expressions during *C. elegans* embryogenesis (*14*).

**Fig. 3.**
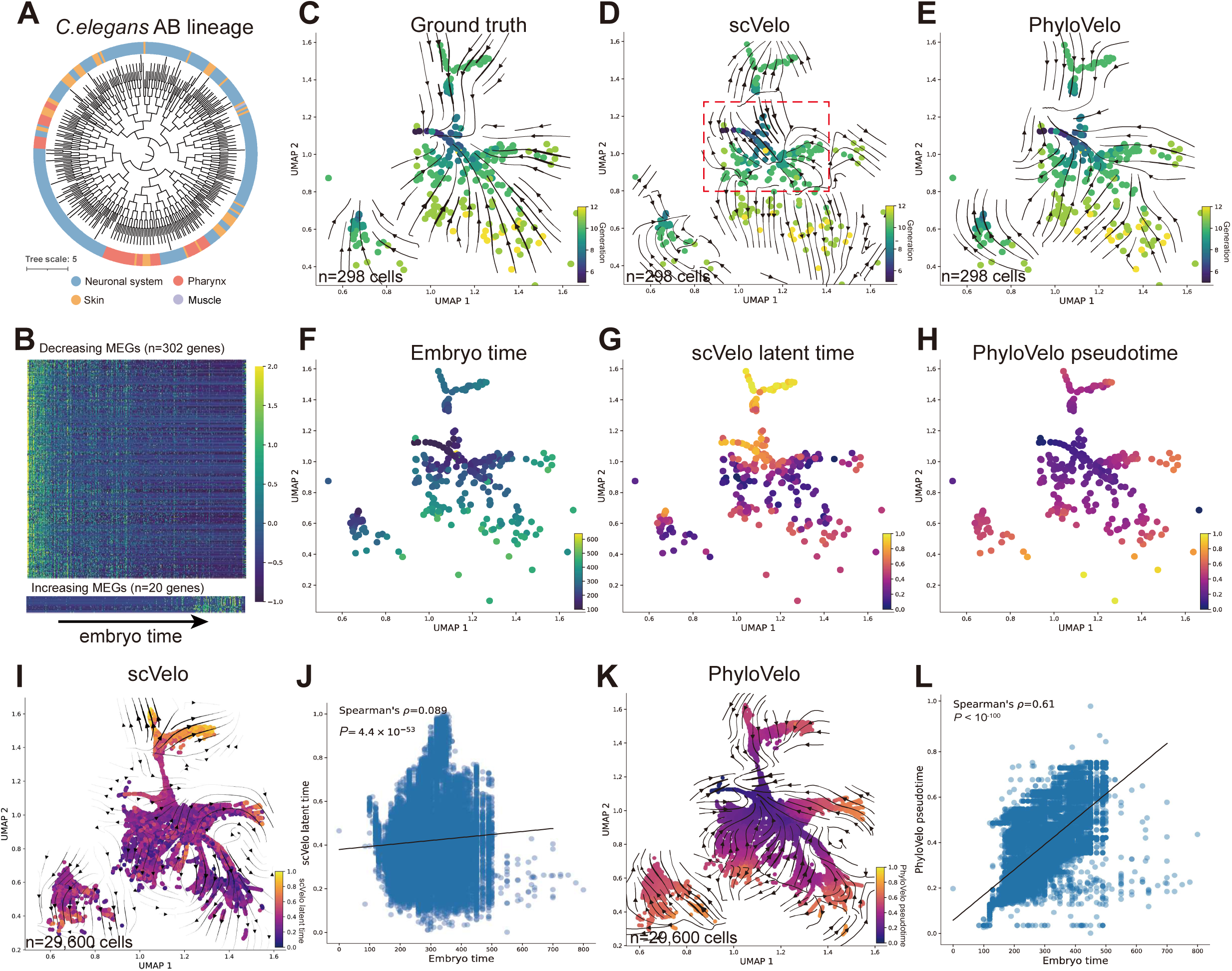
PhyloVelo reconstructs the embryonic differentiation trajectories of *C. elegans*. (**A**) Phylogenetic tree of the *C. elegans* AB lineage. (**B**) Heatmap showing the expressions (z-score normalized) of MEGs along *C. elegans* embryo time. (**C**) The ground-truth velocity fields where vectors are superimposed on the cells that point to each of their immediate parental cells on the Uniform Manifold Approximation and Projection (UMAP) plot. (**D-E**) The velocity fields estimated by scVelo (dynamical mode) (**D**) or PhyloVelo (**E**). Dash square indicates the early embryonic lineages where RNA velocity gave erroneous estimations on the fate directions. (**F**) *C. elegans* embryo time as Packer *et al. (14)*. (**G**) scVelo latent time. (**H**) PhyloVelo pseudotime. (**I**) RNA velocity fields for all 29,600 AB lineage cells. Colors are labeled by scVelo latent time. (**J**) The correlation between scVelo latent time and embryo time for all AB lineage cells. (**K**) PhyloVelo velocity fields for all 29,600 AB lineage cells, estimated by the phylogenetic velocity of MEGs in a single embryo (n=298 cells). Cell colors are labelled by PhyloVelo pseudotime. (**L**) The correlation between PhyloVelo pseudotime and embryo time for all AB lineage cells.

To generate a ground-truth velocity field, each cell was assigned a vector that points to its immediate parental cell on the UMAP plot, which together tracks the cell lineages back to the earliest cells in development (**Fig. 3C**). Therefore, by comparing PhyloVelo with the groundtruth velocity and RNA velocity fields, we were able to evaluate the accuracy of our method. The UMAP embedding clearly reflected the cellular trajectories along cell divisions (**Fig. 3C-E**) or embryo time (**Fig. 3F**). Surprisingly, we found that RNA velocity using scVelo (dynamical mode was used whenever mentioned) failed to recover the expected trajectories with single pseudo-embryo data (**Fig. 3D**), where the scVelo latent time (**Fig. 3G**) was even negatively correlated with the real embryo time in early development before ~300 minutes (**Fig. 3D, Fig. S7A**). In contrast, the directions of phylogenetic velocities reflected the actual development orders (**Fig. 3E**), where the PhyloVelo pseudotime (**Fig. 3H**) was strongly correlated with actual embryo time (Spearman’s ***ρ*** =0.65, *P*=1.4×10^-36^, **Fig. S7B**). Other single pseudo-embryo data also showed similar results (**Fig. S8**).

The RNA velocity fields estimated by all 29,600 AB lineage cells from multiple embryos were improved (**Fig. 3I-J**), suggesting that RNA velocity estimations had been hindered by a small cell number. Remarkably, the phylogenetic velocities of 322 MEGs estimated from single pseudo-embryo data (~300 cells) can be used to correctly estimate the velocity fields for all 29,600 AB lineage cells, even though their lineage trees were not utilized (**Fig. 3K-L**). In fact, the phylogenetic velocities of MEGs were even applicable for non-AB lineage cells, where a convergent trajectory for the first row of head body wall muscle (BWM) and all other BWMs (including C, D and MS lineages) can be identified (**Fig. S9**). These results revealed a general transcriptomic clock during *C. elegans* embryogenesis, and also suggest that compiling a reference phylogenetic velocity of MEGs will greatly facilitate the applications of PhyloVelo to conventional scRNA-seq data where lineage data is unavailable. In summary, the benchmarking on comprehensive simulations and *C. elegans* embryos demonstrated the high robustness of PhyloVelo to recover complex developmental trajectories with phylogeny-resolved scRNA-seq data even with relatively small cell number.

### PhyloVelo resolves multiple-rate kinetics during erythroid differentiation

We next applied PhyloVelo to a CRISPR/Cas9-based lineage tracing dataset from mouse early embryos (E8.0 or E8.5), described by Chan *et al. (32*). This study provided both lineage information and scRNA-seq data via CRISPR lineage tracing of mouse fertilization through gastrulation. By analyzing two embryos with the largest cell number (n=19,017 and 10,800 cells for embryo 2 and 3, respectively), we have identified 469 and 418 MEGs, respectively at whole embryo level (**Fig. S10, Table S1**). Notably, about 40% of MEGs in each embryo were overlapped and the phylogenetic velocities of these overlapped MEGs were strongly correlated (Spearman’s ***ρ***=0.61, *P*=3.2×10^-20^, **Fig. S10**). This indicated the high robustness of our method for identifying MEGs with CRISPR-based lineage tracing data.

To compare the performance of PhyloVelo and RNA velocity, we focused on the erythroid lineage because of its well-defined differentiation trajectory during mouse gastrulation (*46*). Embryo 3 (E8.5) had the largest cell number in erythroid development (n=2,419 cells), thus was selected for this analysis (**Fig. 4A**). RNA velocity failed to identify hematopoietic/endothelial progenitors as the earliest cell types (**Fig. 4B-C**). In addition, the fraction of varying cell types only changed slightly along the scVelo latent time (**Fig. 4D**). In contrast, PhyloVelo correctly predicted the expected trajectory from hematopoietic/endothelial/primitive blood progenitors to primitive blood early/late based on the velocity fields and pseudotime (**Fig. 4E-G**). Importantly, by applying CellRank (*20*), a computational method for identifying the driver genes of cell-fate transitions, with the input of the PhyloVelo pseudotime, we were able to identify the known genes underlying erythroid maturation (e.g. *Alas2, Bpgm, Car2, Slc4a1, Hemgn*, **Fig. S11**).

**Fig. 4.**
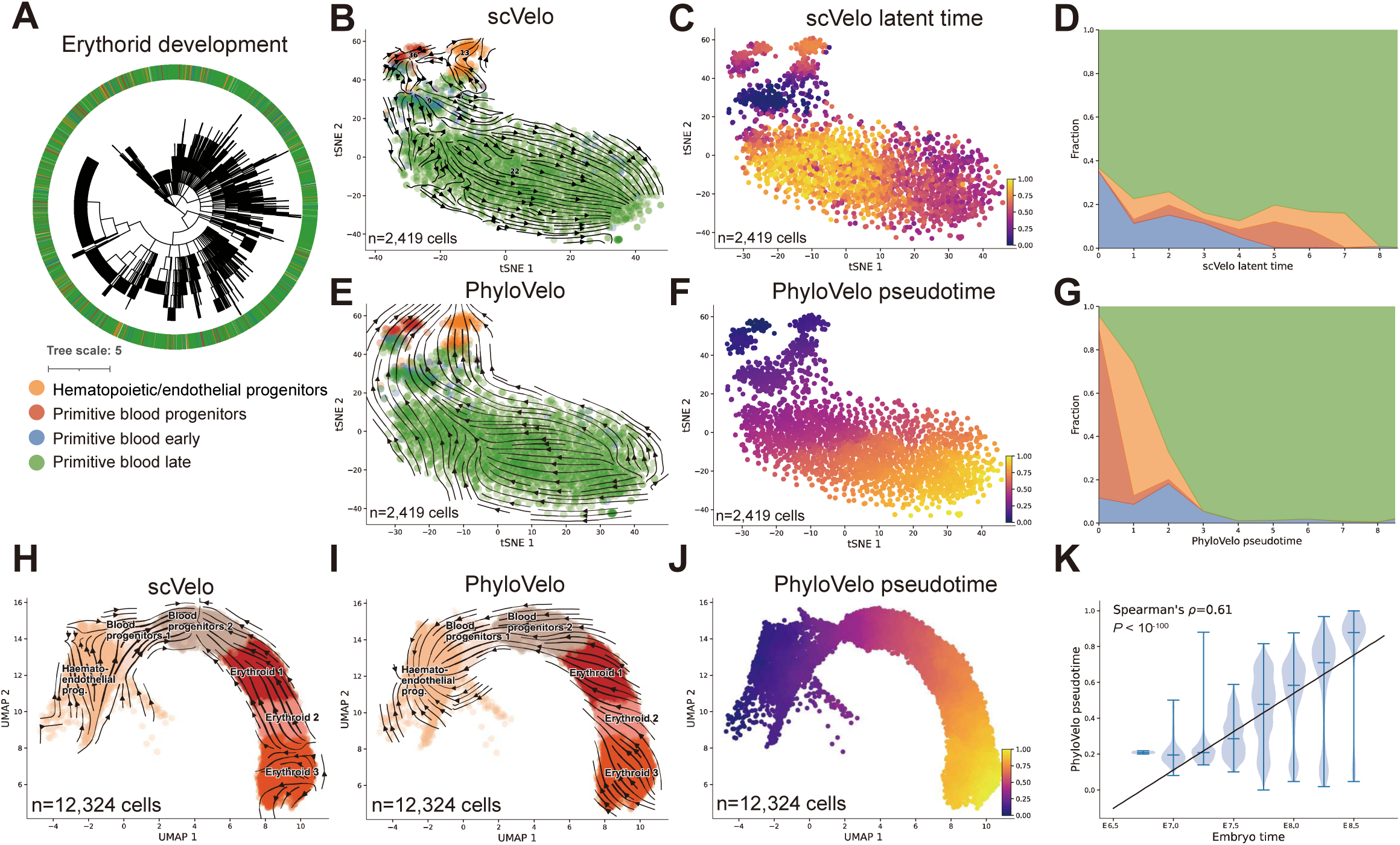
PhyloVelo reconstructs the cellular trajectory of mouse erythroid maturation. (**A**) Phylogenetic tree of the 2,419 erythroid differentiation-related cells (embryo 3, E8.5) in Chan *et al*. dataset *(32)*. (**B-C**) RNA velocity fields (scVelo - dynamical mode) and the latent time of mouse erythroid development. (**D**) Muller plot showing the fractions of four cell types that change over scVelo latent time. (**E-F**) PhyloVelo velocity fields and the pseudotime of mouse erythroid development. (**G**) Muller plot showing the fractions of four cell types that change over PhyloVelo pseudotime. (**H**) Erroneous estimations of RNA velocity fields on erythroid maturation because of multiple rate kinetics (MURK). Data were from Pijuan-Sala *et al. (19)*. (**I**) PhyloVelo velocity fields of erythroid maturation for Pijuan-Sala *et al*. dataset while using the MEGs identified from Chan *et al*. dataset. (**J**) PhyloVelo pseudotime of erythroid maturation in Pijuan-Sala *et al*. dataset. (**K**) The correlation between PhyloVelo pseudotime and mouse embryo time.

Studies have shown that multiple-rate kinetics (MURK) of RNA dynamics violates the model assumptions of RNA velocity, which might lead to erroneous velocity estimations (*24, 25*). Erythroid development is a salient example, where due to MURK, the directions of RNA velocity were even reversed from the expected trajectory of erythroid maturation (*19, 24*) (**Fig. 4H**). Remarkably, using the phylogenetic velocities of the 264 MEGs in erythroid development estimated from single embryo of the Chan *et al.* dataset (*32*) (**Fig. S12, Table S1**), PhyloVelo accurately predicted the expected erythroid trajectory in the scRNA-seq data of temporal mouse embryos (E6.5-8.5) from Pijuan-Sala *et al.* (*19*), despite the lineage tree was not being available (**Fig. 4I**). The PhyloVelo pseudotime was also strongly correlated with mouse embryo time (**Fig. 4J-K**). These data demonstrated that PhyloVelo can circumvent the MURK issue of RNA velocity and the MEGs identified from one dataset can be also applied to independent datasets, even when phylogenetic information is not available.

### PhyloVelo reconstructs a dedifferentiation trajectory in lung tumor evolution

We next applied PhyloVelo to a CRISPR/Cas9-based lineage tracing dataset in a genetically-engineered mouse model (GEMM) of lung adenocarcinoma (*Kras*^LSL-G12D/+^;*Trp53*^fl/fl^, or KP model), described by Yang *et al.* (*47*). Cancer GEMMs allow one to study tumor evolutionary trajectory in its native microenvironment. Two primary tumors from KP mice (3726_NT_T1 and 3435_NT_T1) were selected because of their relatively high-resolution lineage trees and composition of diverse cell types (including AT2-like, AT1-like, Gastric-like, High plasticity, Lung-mixed, Endoderm-like, Early EMT (epithelial–mesenchymal transition)-1, etc.) (**Fig. 5A, Fig. S13A**). In total, 339 and 349 MEGs were identified from these two tumors, respectively (**Fig. S14, Table S1**). RNA velocity by scVelo performed reasonably well in 3435_NT_T1 to recapitulate the expected trajectory from AT2-like to High plasticity, and to Lung-mixed cells (**Fig. S13B**), whereas no clear trajectory was inferred in 3726_NT_T1 by scVelo (**Fig. 5B**). In contrast, in both tumors PhyloVelo identified AT2-like cells as the cell-of-origin of KP lung adenocarcinoma and also recovered the expected trajectory from AT2-like to lung-mixed or Early EMT (**Fig. 5C, Fig. S13C**). In 3726_NT_T1, two trajectories appeared to coexist, namely 1) AT2-like > lung-mixed> Early EMT and 2) AT2-like > Endoderm-like > Early EMT (**Fig. 5C**), thus recapitulating the findings in the original study (*47*).

**Fig. 5.**
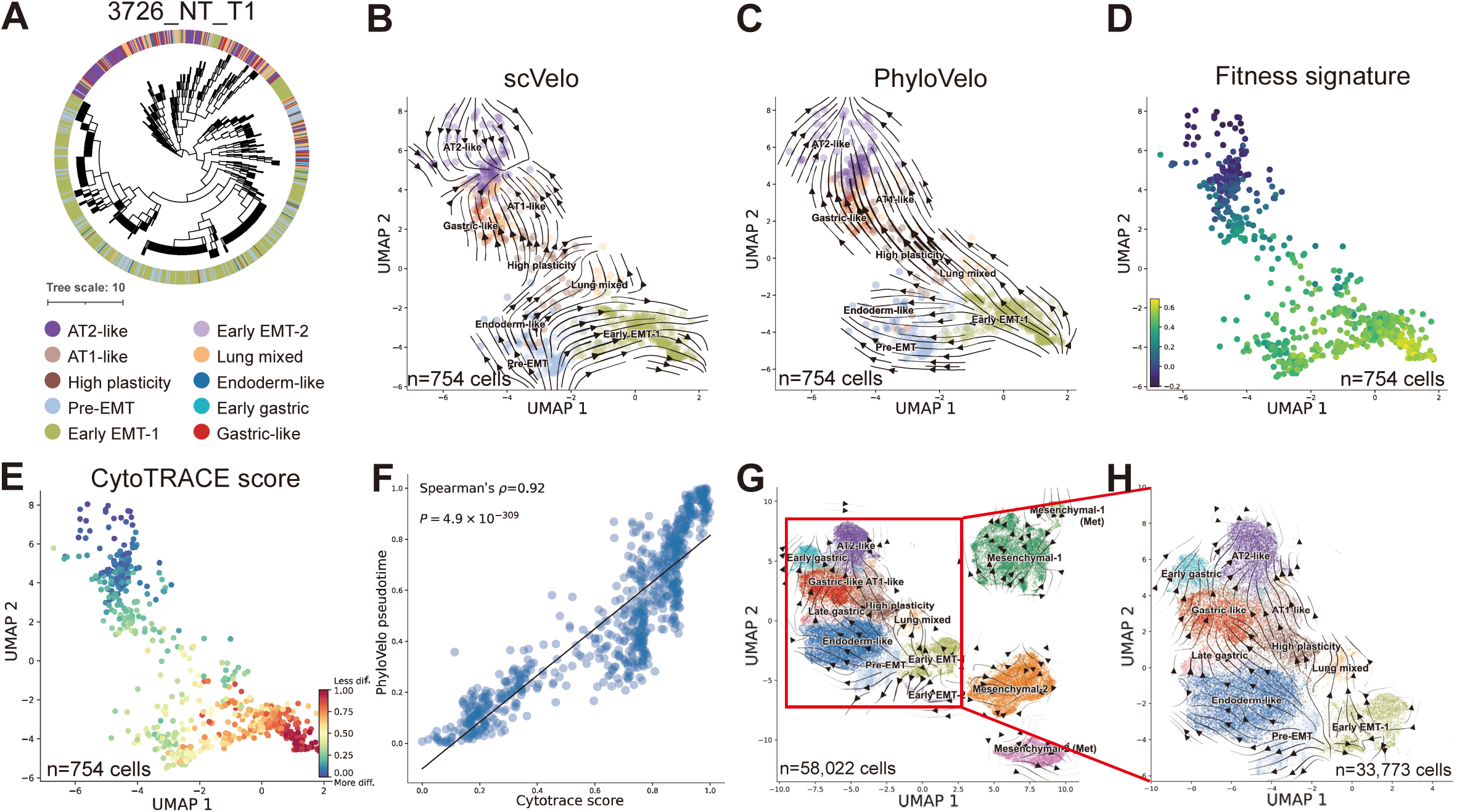
PhyloVelo identifies a dedifferentiation trajectory in lung tumor evolution. (**A**) Phylogenetic tree of 754 cells from a KP-mouse primary lung tumor, 3726_NT_T1, in Yang *et al*. dataset *(47)*. The scRNA-seq data, cell type annotations, and lineage trees were obtained from the original study. (**B**) RNA velocity fields (scVelo - dynamical mode) for 3726_NT_T1. (**C**) PhyloVelo velocity fields for 3726_NT_T1. (**D**) Fitness signatures of single cells for 3726_NT_T1, as defined by Yang *et al*. (**E**) CytoTRACE score of single cells for 3726_NT_T1. (**F**) The correlation between PhyloVelo pseudotime and CytoTRACE scores. (**G**) PhyloVelo velocity fields for all 58,022 single cells from pooled KP primary lung tumors, estimated by the MEGs identified from 3726_NT_T1. (**H**) PhyloVelo velocity fields for the cell types that existed in 3726_NT_T1.

Yang *et al.* (*47*) defined a single-cell fitness signature (**Fig. 5D, Fig. S13D**) by a specific gene module whose expression is associated with the proliferating fitness of the corresponding cell. While no overt association between scVelo latent time and the fitness signatures was found (**Fig. 5B, Fig. 5D, Fig. S13E**), the PhyloVelo pseudotime showed a strong correlation with the fitness signature in both tumors (3726_NT_T1, Spearman’s ***ρ***=0.86, *P*=1.2×10^-218^; 3435_NT_T1, Spearman’s ***ρ***=0.83, *P*=4.0×10^-300^, **Fig. 5C, Fig. 5D, Fig. S13F**). This indicates an intrinsic link between our measure of phylogenetic velocity and the cell fitness. Interestingly, CytoTRACE (*10*), a computational algorithm to predict the cellular differentiation states with scRNA-seq data, revealed a drastic increase of the expressed gene counts in the tumor evolution (**Fig. 5E**, **Fig. S13G**), which was in consistent with a dedifferentiation model. This also indicated that although gene expression diversity is a key feature of the developmental potential (*10*), the directions of cell-state transitions are highly context-dependent and the cellular trajectories in normal differentiation and disease progression can be completely reversed.

We further demonstrated the phylogenetic velocity of MEGs identified from only one pilot tumor (3726_NT_T1, **Fig. S14**) allowed for the robust inference of velocity fields for other independent KP tumors where the lineage trees were not utilized (n=58,022 cells) (**Fig. 5G-H**). In fact, the inferred and expected trajectories were highly consistent for the cell types that existed in 3726_NT_T1 but not for the novel cell types such as Mesenchymal 1 and 2 (**Fig. 5G-H**). Interestingly, the phenomenon that two trajectories coexist as in 3726_NT_T1 (**Fig. 5D**) was more evident on the pooled PhyloVelo velocity fields (**Fig. 5H**). These results demonstrated the generality of transcriptomic clocks across the KP tumors, but also implied that a large single-cell lineage tree spanning numerous cell types must be reconstructed in order to identify more ubiquitously clock-like MEGs.

### Ribosomal genes underlie the cellular transcriptomic clock

To systematically investigate the potential functions of MEGs across tissues and organisms, we analyzed three additional CRISPR-based lineage tracing datasets that were derived from mouse or human cell lines, including pancreatic cancer KPCY (*48*), lung cancer A549 (*49*) and normal epithelial cells HEK293T (*50*). The KPCY and A549 cells were sampled from *in vivo* mouse xenograft model, while the HEK293T cells were from *in vitro* culture. Interestingly, although these cell lines are known to be non-differentiating, continuous cell-state transitions were remarkable according to the PhyloVelo velocity fields (**Fig. 6A-C, Fig. S15-17**). For instance, PhyloVelo recovered a dynamic EMT trajectory during the metastatic progression of KPCY and A549 mouse xenografts (**Fig. 6A-B, Fig. S15-16**), which again outperformed RNA velocity (**Fig. S15-16**). Even for *in vitro* culture of HEK293T cells, both PhyloVelo and scVelo revealed continuous state transitions (**Fig. 6C, Fig. S17**). Interestingly, the CytoTRACE “stemness” scores were strongly and positively associated with PhyloVelo pseudotime in mouse xenografts of A549 cells (**Fig. S18**), also indicating a dedifferentiation process during the tumor progression of mouse xenograft.

**Fig. 6.**
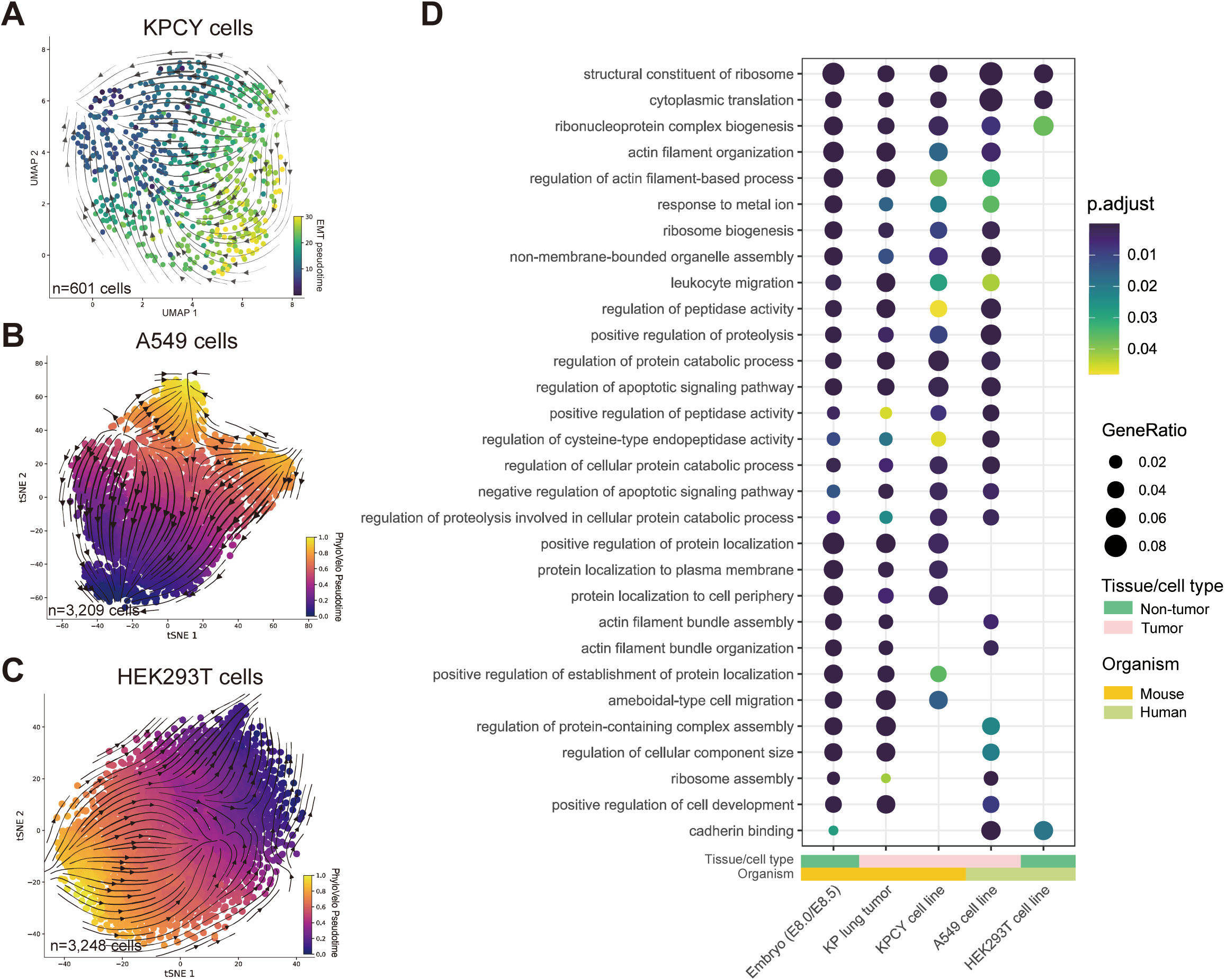
MEGs are enriched in ribosome-mediated processes across tissues and organisms. (**A**) The EMT trajectory of KPCY cells in mouse xenograft model reconstructed by PhyloVelo. Cell colors are labelled by EMT pseudotime. (**B-C**) Continuous state transitions in mouse xenograft model of A549 cells (**B**) and *in-vitro* clonal culture of HEK293T cells (**C**), respectively. The largest clonal population of A549 cells, lg1, is shown here. Cell colors are labelled by PhyloVelo pseudotime. (**D**) Gene ontology (GO) enrichment of MEGs identified across tissues and organisms. The top and most commonly shared 30 biological processes are shown.

We found the MEGs identified across organisms (mouse and human) and tissue or cell types (embryo, primary tumor, cancer and normal cell lines) were significantly overlapped (**Fig. S19, Table S1**). Remarkably, the ribosome machinery was strongly enriched across the tissues and organisms, including translation, ribonucleoprotein complex biogenesis, ribosome biogenesis, and others (**Fig. 6D**). To rule out the possibility that ribosomal genes were identified because of their high expression heterogeneity amongst the cells, we further performed permutation analysis where the phylogenetic distances were randomly shuffled and assigned to the cells. Followed by the PhyloVelo inference procedures, “pseudo-MEGs” can be identified if their latent expressions also showed significant association with the randomly assigned phylogenetic distances. Here, although many “pseudo-MEGs” can be identified (**Fig. S20A**), they only showed weak associations (most *p* values were around 0.05) with the phylogenetic distances. Importantly, no significant enrichment in ribosome-mediated processes was found (**Fig. S20B**). These results strongly suggest that many ribosomal genes genuinely follow the clock-like expression dynamics during cell proliferation and differentiation.

## Discussion

Defining the correct directions of cell-fate transitions is crucial for unraveling the genetic or epigenetic factors that drive lineage specification in diverse biological contexts (*20*). Although RNA velocity and its improvements (*22, 23, 51, 52*) are powerful approaches for quantifying cellular transitions from single-cell transcriptomic data, an accurate estimation of the velocity fields is still challenging because of the highly dynamic RNA transcription rates (*23–25*) and biased capture of intron regions by droplet-based scRNA-seq (*53*). The fundamental objective of our PhyloVelo algorithm is the same as RNA velocity – to extrapolate the gene expression of single cells to their near future or past states. However, unlike RNA velocity, PhyloVelo quantifies the transcriptomic velocity by measuring the rate of expression changes along a cell division history. Using various single-cell datasets where the coupled lineage trees were available, PhyloVelo not only recovered the expected trajectories more accurately, but also gave more consistent estimates of the velocities relative to RNA velocity, across diverse biological contexts.

Analysis of phylogeny-resolved scRNA-seq datasets across embryo development, normal cell expansion and tumor evolution with PhyloVelo yields new insights into cell-state dynamics. First, in each of the lineage tracing datasets, we have identified 200-500 MEGs, suggesting a considerable number of genes show directional expression trajectories along cell divisions. Interestingly, the MEGs across tissues and organisms had highly similar functions in translation and ribosome biogenesis, which actually are canonical hallmarks of cell cycle and proliferation (*54*). Interestingly, ribosomal genes have also been found to exhibit high expression heterogeneity in different cell types, developmental stages, and even within the same cell type (*55–57*). We speculate that the expression heterogeneity of ribosomal genes might be partially caused by the intrinsic variability of cellular proliferating capacity. Indeed, ribosome biogenesis had been implicated in cancer (*58, 59*). Second, we demonstrated that the phylogenetic velocities of MEGs estimated from one phylogeny-resolved scRNA-seq dataset can be used to infer the lineage trajectory in independent scRNA-seq datasets, in the absence of phylogenetic information. Because obtaining a coupled lineage tree for every scRNA-seq dataset is rather laborious, the transferability of MEGs greatly facilitates the application of PhyloVelo to conventional scRNA-seq datasets. Therefore, future efforts should focus on identifying and compiling a ubiquitous panel of MEGs and estimating their phylogenetic velocities at the whole-organism or whole-organ level, and evaluating them with PhyloVelo in different tissues.

Despite the rapid development of CRISPR lineage tracing methods (*4, 29, 60*), building high-precision lineage trees with single-cell resolution is still challenging because of the small number of Cas9 target sites (typically<50), rapid saturation, frequent inter-site deletions, and other factors (*61–63*). This requires a more reliable lineage tracing method that has a larger lineage-labeling space and more stable mutagenesis strategy. We recently developed a base editor-based lineage tracing method, called SMALT (*41*) (Substitution Mutation-Aided Lineage-Tracing), which leverages a genetically-evolved activation-induced cytidine deaminase (AID) to specifically target a 3k synthetic DNA barcode and induce C to T mutations on it with high efficacy. The lineage tree reconstructed by SMALT achieved nearly single-cell resolution and over 80% statistically bootstrapping support (*41*). Importantly, AID-induced C to T mutations only occur during DNA replication and thus are mitosis dependent. Thus, the number of mutations in this system enables more reliable measurement of the number of cell divisions, which is more suitable for PhyloVelo inference. We envision the combination of SMALT lineage tracing and single-cell transcriptomics will greatly empower the application of PhyloVelo to resolve complex lineage dynamics in more diverse biological contexts, such as genetic perturbation or disease progression.

In summary, we provide a novel theoretical framework and a powerful method to quantify cell-fate transitions by exploiting both single-cell phylogenetic and transcriptomic information. With the rapid development of single-cell lineage tracing technologies and emergence of large-scale lineage-traced genomic data, we envision our method will greatly facilitate the lineage analysis for complex cellular processes and the discovery of the cell-fate determinants in diverse organisms, tissues, and diseases.

## Methods

### The mathematical framework of PhyloVelo

The dynamics of the latent expression *z* for each gene on a phylogeny *T* was assumed to follow a diffusion process (also known as the stochastic differential equation, SDE), which varies along cell divisions:

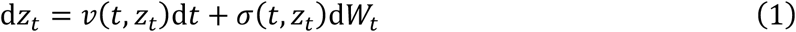

Here, *W_t_* is a standard Brownian motion. In our model, we hypothesized that there is a group of genes *G_m_* whose drift coefficient *v*(*t, z_t_*) and diffusion coefficient *σ*(*t, z_t_*) are independent of both *t* and *z_t_*, thus *v*(*t, z_t_*) = *v* and *σ*(*t, z_t_*) = *σ*. We called them monotonically expressed genes (MEGs). For this type of genes, the dynamics of its latent expression *z* is thus formulated as:

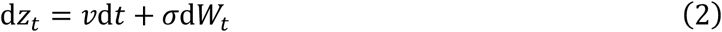

and its expectation is given by:

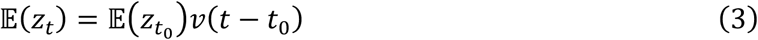

For the observed scRNA-seq measurement *x* (read or UMI count), we assumed that it is sampled from the negative binomial (NB) distribution or the zero-inflated negative binomial (ZINB) distribution:

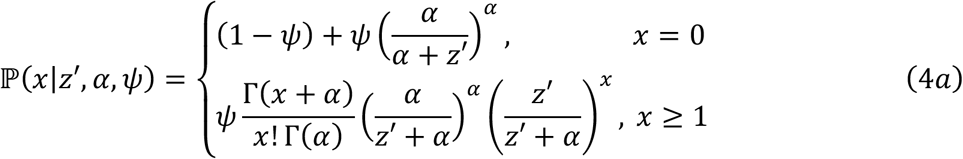

where *z*′ is the exponential function of latent expression *z, α* is the scale parameter, and *ψ* is the zero-inflation parameter. The expectation of the distribution is *z*′*ψ*. For the negative binomial distribution, *ψ* = 1. We used the likelihood ratio test to verify zero inflation for each gene (**Supplementary Text**).

For the scRNA-seq data after normalization (e.g. using *scanpy.pp.normalize_per_cell* and *scanpy.pp.log1p*), we also provided a Gaussian model of latent expression for normalized data:

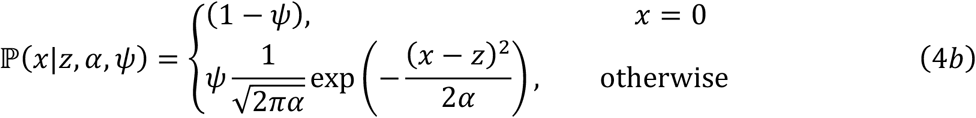

To estimate the latent expression *z*, we used the maximum a posteriori probability (MAP) estimate. For the ZINB model using raw UMI count data, we estimated *z*′ and then took logarithm to get the estimated latent expression *z*:

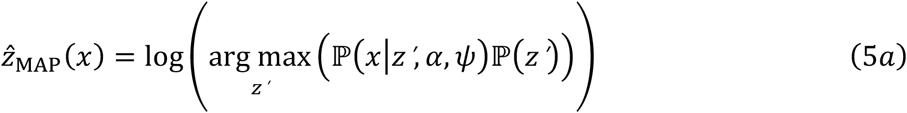

For the Gaussian model using normalized UMI count data, we directly performed the MAP estimate of the latent expression *z*:

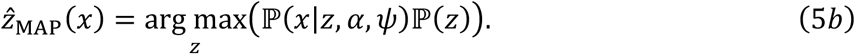

For a MEG *g*, its drift coefficient can be estimated as:

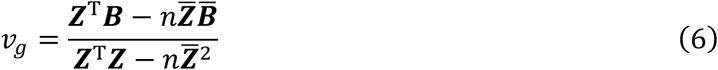

Here 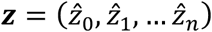 represents the estimated latent expressions and ***B*** = (*b*_1_, *b*_2_, …, *b_n_*) the phylogenetic distances from terminal cells to the root (*n* is the cell number in a dataset). The drift coefficients of all *G* MEGs in a dataset *v* = (*v*_1_, *v*_2_,⋯, *v_G_*) were thus referred to be as phylogenetic velocity.

### Simulation of phylogeny-resolved scRNA-seq data

To generate simultaneous single-cell phylogeny and transcriptome data *in silico*, a lineage-embedded scRNA-seq data simulator, PROSSTT (*40*), was modified to account for dividing cell populations, so that the whole cell division history initiated from a single cell can be recorded. The simulation consisted of three parts:

1. Simulate a cell dividing and differentiating process using the Gillespie algorithm to obtain the cell division history;
2. Given a cell differentiation model (linear, bifurcated or convergent), use the diffusion process to generate gene expression programs;
3. Assign the gene expression programs onto the cell division history in order to obtain the read count data for each gene in each cell.

#### Simulating cell division history and mutation-based phylogeny

We used a continuoustime Markov process to simulate cell growth and differentiation. In particular, each cell type *i* has a specific division rate *p_i_*(*t*) and differentiation rate *q_ij_*(*t*), given as follows:

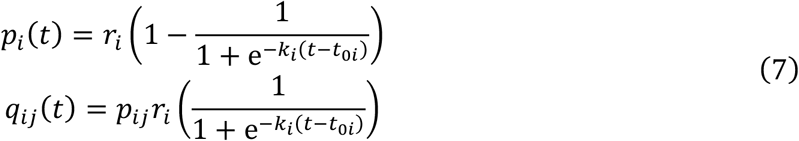

where *r_i_, k_i_* and *t*_0*i*_ are the cell-type specific parameters and *p_ij_* is the probability of cell type *i* differentiating into cell type *j*, ∑_*i*≠*j*_ *p_ij_* = 1, *p_ii_* = 0.

Now, we can simulate the cell growth process using the Gillespie algorithm (*64*). Each simulation ended when the population size reached 10,000 cells. Then, 1,000 cells were randomly sampled to obtain their division history.

To simulate the mutation-based cell phylogeny, we assumed that mutations randomly occurred during each cell division by a Poisson distribution:

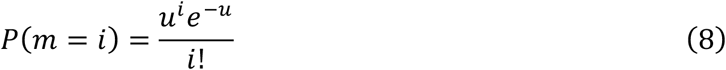

where *u* is the mean mutation rate per cell division. Different mutation rates (*u* =0.1, 0.3, or 1) were used. After obtaining the cell-mutation matrix, we used three different algorithms to reconstruct the phylogenetic tree, respectively, namely *Maximum Likelihood* (using IQ-TREE 2 (*65*)), *Neighbor-Joining* (using R package ape 5.6-2 (*66*)) and *Maximum Parsimony* (using MEGA X (*67*)).

#### Simulating scRNA-seq data

We first simulated the latent expression process of genes. For each gene, we randomly generated its initial expression *μ*_0_, drift coefficient *v*, and variance *σ*^2^, and then simulated gene-specific diffusion process as follows:

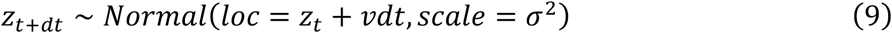

When cells differentiated at time *t_d_*, for MEGs, their gene expressions remained unchanged as the same with the values in the previous process. For a non-MEG, its drift coefficient and variance were regenerated randomly and the value of expression was reset to *z_t_d__*. We called the diffusion process *z_t_* as the gene expression program. For each gene, the initial value of the expression program *z*_0_ was randomly drawn from a gamma distribution *z*_0_ ~ Γ(0.5,20), the drift coefficient *v* was drawn from a normal distribution *v* ~ Normal(0,1), and the diffusion coefficient *σ* was drawn from a truncated normal distribution 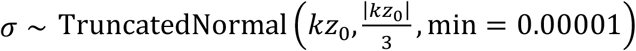. In all simulations, we set the drift coefficient of each gene expression program to change with probability 0.4 when cell differentiation happens, ultimately resulting in 10-15% of genes with unchanged expression programs upon cell differentiations, thus behaving as MEGs. The other 85-90% of genes will change dynamically with cell differentiations and thus behave as non-MEGs. Each cell was assumed to have 2,000 expressed genes, thus including 200-300 MEGs in total in each simulated dataset.

After generating the latent expression process of all genes, in order to simulate the variations introduced in real scRNA-seq experiments, the NB (*68*) or ZINB distribution (*69*) was used to obtain read/UMI count *x*:

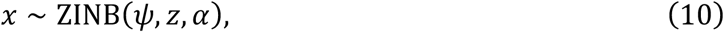

where *ψ* is the zero-inflation parameter (*ψ* = 1 for negative binomial model), *α* is the scale parameter and the expectation of the distribution is *ψz.*

#### Assigning the gene expression programs to the cell division history

Having each gene expression program (*z*_1_, *z*_2_, ⋯, *z_G_*) and phylogeny 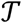, we traversed all nodes 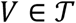 and assigned the latent expression programs *z*_1_(*d*), ⋯ *z_G_*(*d*) to the nodes with branch length of *d.* For each gene *g*, a random number obeying the distribution *z_g_*(*d*) was drawn as the latent expression of that gene. By traversing all the genes and cells, the latent expression matrix can be obtained and denoted as *Z.* We also simulated a Gaussian noise *ε* to be imposed on *Z*, thus the latent expression matrix would be updated as *Z* + *ε*. Finally, given the zero-inflation factor *ψ* and the scale parameter *α*, the expression matrix *X* of the simulated data can be obtained by random sampling according to Equation (10).

### Projection of phylogenetic velocities into the low dimensional embedding

Inspired by RNA velocity (*22*) and scVelo (*23*), phylogenetic velocities were projected into the dimensionality reduction embedding using the following method.

Given the phylogenetic velocity *v*, the gene expression matrix ***X**_*n*×*m*_ = (***x***_1_, ***x***_2_,⋯, ***x***_*n*_)^*T*^, and an embedding with *n* positions of cells ***S*** = (***s***_0_, ***s***_2_,⋯, ***s**_n_*), where *n* is the number of cells, *m* is the total number of genes, ***x**_i_*, = (*x*_*i*1_, *x*_*i*2_, ⋯, x_im_*) For cell *i*, we first calculated the difference between ***x**_i_* and other cell expression vectors ***x***_*j*(*j*≠*i*)_ as follows:

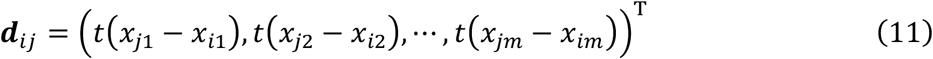

where 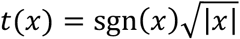. We then used an exponential kernel transformation as the weight:

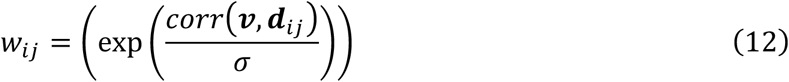

where *corr* represents Pearson correlation coefficient. Finally, the weighted sum of the base vectors was used to project the velocity into embedding:

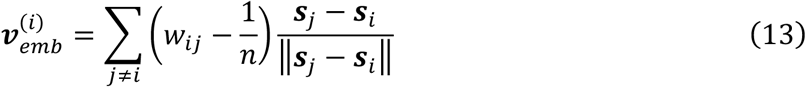

### Inference of PhyloVelo pseudotime

To infer the PhyloVelo pseudotime of each cell, we first constructed a minimum spanning tree based on the distance of cellular states on tSNE/UMAP embedding using Prim’s algorithm (*70*). Thus, we can obtain a subset of the edges *ℇ* connecting all cells together with a minimum possible total distance. We then chose any cell *c*_0_ as the starting point and set its pseudotime *pt*_*c*_0__ = 0. For any other cells 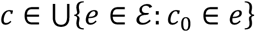, we calculated its pseudotime using the following equation:

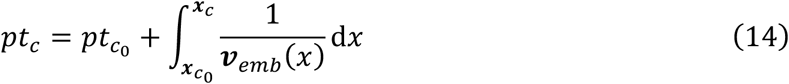

where ***x**_c_* is the coordinate of cell *c* in the embedding space and ***v**_emb_*(*x*) is the phylogenetic velocity in the embedding space and varies with its coordinates.

To simplify the calculation, we replaced the velocity in this path with the average velocity of ***v***(*c*) and ***v***(*c*_0_), denoted by ***v**_a_*, and used the line segments ***l***_*c,c*_0__ to approximate the path. Hence, we have:

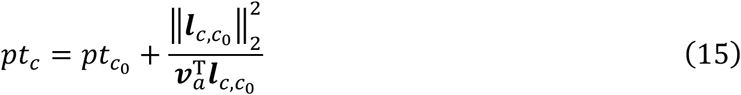

Following the path generated from the minimum spanning tree, we can estimate the pseudotime of all cells and finally normalize to [0,1].

### Analysis of real phylogeny-resolved scRNA-seq datasets

#### Datasets and pre-processing

We have applied PhyloVelo to six real phylogeny-resolved scRNA-seq datasets that are publicly available through online sources (see **Data availability**). These included *C. elegans* (*14*), mouse embryos (*32*), GEMM of lung adenocarcinoma (*47*), mouse xenograft models of pancreatic cancer cell line KPCY (*48*) and lung cancer cell line A549 (*49*), and *in-intro* culture of human kidney cell line HEK293T (*50*). The embryonic lineage tree of *C. elegans* is entirely known and was obtained from http://dulab.genetics.ac.cn/TF-atlas/Cell.html, while the CRISPR-based lineage trees in other five datasets were obtained from the original studies. In the *C. elegans* dataset, because multiple synchronous embryos were pooled for the scRNA-seq experiment, many nodes in the lineage tree have been sampled multiple times. Thus, only one random cell was chosen to represent the corresponding node, while these non-repetitive cells (~300 cells) from one lineage tree constituted a “pseudo-embryo”. For the scRNA-seq data of *C. elegans (14*) and mouse lung adenocarcinoma (*47*), the coordinates of tSNE or UMAP from the original studies were used. For the scRNA-seq data of mouse embryos (*32*), cell lines KPCY (*48*), A549 (*49*) and HEK293T (*50*), the dimensionality reduction and tSNE visualization were performed using Scanpy (*71*) following the recommended data processing procedures and parameters as https://scanpy-tutorials.readthedocs.io/en/latest/. In each dataset, the genes with total count < 20 were filtered out.

#### Applying PhyloVelo

For *C. elegans*, whose embryonic cell division history is entirely known, the cell generation time was used to denote the phylogenetic distance. For the other five CRISPR/Cas9 lineage tracing datasets (*32, 47–50*), the phylogenetic distance on a lineage tree corresponds to the expected number of Cas9 cutting scars on the evolving barcodes. To estimate the latent gene expressions, for *C. elegans*, the ZINB model was used to analyze the raw UMI count data because of the high-quality lineage tree. For the CRISPR/Cas9-based lineage tracing datasets, the Gaussian model was used on the post-normalized data where *normalize_per_cell(*) and *log1p* by *Scanpy (71*) were applied to the raw UMI counts. To prioritize the high-confident candidates of MEGs and speed up the computation, rather than estimating the latent expression for all genes, we firstly searched for candidate MEGs by directly analyzing the correlation between the normalized UMI counts and the phylogenetic distances of single cells. The top 5% of genes with the highest Spearman’s correlations were selected and proceeded for latent expression estimations as well as MEGs identification. Final MEGs were identified by their significant association (Spearman’s correlation, *P*<0.05) between the latent expressions and the phylogenetic distances from terminal nodes to the root of tree. The phylogenetic velocity was computed independently for each MEG. To project the phylogenetic velocity into the dimensionality reduction embedding, we built a k-nearest neighbor (kNN) graph (k=15 for *C. elegans* dataset while it was chosen by approximate to one third of total number of cells for the CRISPR lineage tracing datasets). The kNN graph was based on the Euclidean distance as the base vector and was used to estimate the coordinates of velocity embedding, as the projection of RNA velocity (*22, 23*).

#### Applying scVelo

The spliced and unspliced read counts were obtained by running velocyto (v0.6) (*22*) on the bam files from the output of CellRanger (6.0.2) on the raw sequence reads. To estimate RNA velocity, scVelo (version 0.2.4) (*23*) and the dynamical mode were used following the recommended data processing procedures as https://scvelo.readthedocs.io/VelocityBasics/.

#### Gene ontology (GO) enrichment analysis

GO enrichment analysis was performed using clusterProfiler v4.4.4 (*72*). The cutoff for *p* value and *q* value were set to 0.05 and 0.25, respectively. After excluding Cellular Components (CC) terms, top 10 GO terms were retained for downstream analyses. Subsequently, top 10 GO terms of each sample were merged and these terms were sorted by their total occurrence and minimal *q* value across samples. The GO terms enriched in more than two datasets were visualized using ggplot2 v3.3.6.

### Transferring phylogenetic velocities to independent scRNA-seq datasets

To evaluate whether the phylogenetic velocities of MEGs estimated from one phylogeny-resolved scRNA-seq dataset are sufficiently robust to infer the velocity fields in independent datasets in the absence of phylogenetic information, three datasets were analyzed including *C. elegans* (*14*), mouse erythroid development (*19, 32*), and the GEMM of lung adenocarcinoma (*47*). Here, the MEGs and corresponding phylogenetic velocity estimates were directly applied to another scRNA-seq datasets without the lineage tree. For *C. elegans*, we applied the phylogenetic velocities from AB lineage in a single pseudo-embryo (n=298 cells) to all AB lineage cells (n=29,600) in multiple embryos. We also applied them to non-AB lineages that differentiate to body wall muscles (BWM) (including C, D and MS lineages; n=1,992 cells). For mouse erythroid differentiation, we applied the phylogenetic velocities estimated by the erythroid lineage cells in a single embryo (E8.5, n=2,419 cells) in the Chan *et al.* dataset (*32*) to the temporally-sequenced mouse embryos (E6.5-E8.5, n=12,324 cells) in Pijuan-Sala *et al.* dataset (*19*). Finally, for lung adenocarcinoma (*47*), the phylogenetic velocities estimated from one KP primary lung tumor (3726_NT_T1, n=754 cells) were applied to all 58,022 single cells in pooled KP primary lung tumors.

## Data availability

All data analyzed in this article are publicly available through online sources. The annotated data, lineage trees, results and Python implementation are available at https://phylovelo.readthedocs.io/. The raw data for the *C. elegans* dataset can be accessed at the Gene Expression Omnibus database with accession number GSE126954, the CRISPR lineage tracing datasets from the mouse embryos with accession number GSE117542, the mouse primary lung tumors with accession number PRJNA803321 and Zenodo Data (https://zenodo.org/record/5847462#.Yt4-PewRXUI) and mouse pancreatic cancer cell line KPCY with accession number GSE173958 and Mendeley Data (https://doi.org/10.17632/t98pjcd7t6.1), the human lung cancer cell line A549 with accession number GSE161363 and the human kidney cell line HEK293T with accession number PRJNA757179.

## Code availability

PhyloVelo is freely available as Python package at https://github.com/kunwang34/PhyloVelo. Detailed workflows to reproduce figures and results in this paper are written as Jupyter notebook in the repository. The annotated data, lineage trees, results and Python implementation are available at https://phylovelo.readthedocs.io/.

## Acknowledgements

We thank Yuanhua Huang, Jin Xu, Liang Ma, Wanze Chen, and Hu laboratory members for constructive discussions. This work was supported by National Key R&D Program of China (2021YFA1302500 to Z.H.), Guangdong Basic and Applied Basic Research Foundation (2021B1515020042 to Z.H.), National Natural Sciences Foundation of China (11971405 to D.Z., 32270693 to Z.H.) and China Postdoctoral Science Foundation (2021M693303 to Z.L).

## Author contributions

Z.H. and K.W. conceived the concept of phylogenetic velocity. Z.H., K.W., and D.Z. designed the study. K.W. developed the mathematical framework and implemented the software. K.W., Z.H., L.H., Z.L., X.W., Z.Y. analyzed the data. W.Z. and Z.Z. provided constructive suggestions on the model. K.W., Z.H., D.Z., X.H., C.C., interpreted results. Z.H. and K.W. wrote the manuscript with contributions from all co-authors. Z.H., and D.Z. supervised the study.

## Supplementary Materials

Supplementary Text

Figs. S1 to S20

Table S1

