## Supplementary Materials for "Cell division history encodes directional information of fate transitions"

Kun Wang, *et al.*

 (Z.H.)

**This PDF file includes:**

Supplementary Text
Figs. S1 to S20
Table S1

### Supplementary Text

#### Hypothesis testing for zero inflation in scRNA-seq data

To determine whether there is zero inflation for the expressions of a gene amongst the cells, we constructed the likelihood ratio statistic, $LR$, defined by:

$$LR = 2\log\left(\frac{\mathcal{L}(\hat{\theta}_1|\psi \neq 1)}{\mathcal{L}(\hat{\theta}_0|\psi = 1)}\right) \tag{S1}$$

where $\hat{\theta}_1$ and $\hat{\theta}_0$ are the maximum likelihood estimates for the conditions $\psi = 1$ and $\psi \neq 1$, respectively. $\mathcal{L}$ is defined by:

$$\mathcal{L} = \sum_{n=1}^{N} \log(\mathbb{P}(x_i)) \tag{S2}$$

It is easy to prove that:

$$LR \sim \chi^2(3), (N \to +\infty) \tag{S3}$$

Therefore, the rejection region of the hypothesis testing problem is:

$$R^+ = \{LR(\theta) > \chi^2_{1-\alpha}(3)\} \tag{S4}$$

#### Estimation of latent expression $\boldsymbol{z}$

To get the maximum a posteriori estimation (MAP) of $\boldsymbol{z}$ as shown in equation (5a), we took its partial derivative with respect to the exponential function of latent expression, $z'$:

$$\begin{aligned}\frac{\partial \mathbb{P}(x|z',\alpha,\psi)\mathbb{P}(z')}{\partial z'} &\propto \frac{\partial}{\partial z'}\left(\left(\frac{\alpha}{\alpha+z'}\right)^{\alpha}\left(\frac{z'}{\alpha+z'}\right)^{x}\exp\left(-\frac{(z'-z_0')^2}{2\sigma^2}\right)\right) \\ &= -\mathrm{e}^{-\frac{(z\prime-z_0')^2}{2\sigma^2}}\left(\alpha\sigma^2(z'-x)+\alpha z(z'-z_0')+z'^2(z'-z_0')\right) \\ &\quad \cdot \frac{\left(\frac{\alpha}{\alpha+z'}\right)^{\alpha}\left(\frac{z'}{\alpha+z'}\right)^{x-1}}{\sigma^2(\alpha+z')^2}\end{aligned} \tag{S5}$$

Setting equation (S5) to $0$, we have:

$$\begin{aligned} z_1' &= 0, \\ z_2' &\approx 0.26457\Delta_3 - \frac{0.41997\Delta_1}{\Delta_3} + 0.3333(z_0'-\alpha), \\ z_3' &\approx -(0.13228-0.22912i)\Delta_3 - \frac{(0.20999+0.36371i)\Delta_1}{\Delta_3} + 0.3333(z_0'-\alpha), \\ z_4' &\approx -(0.13228+0.22912i)\Delta_3 + \frac{(0.20999-0.36371i)\Delta_1}{\Delta_3} + 0.3333(z_0'-\alpha) \end{aligned} \tag{S6}$$

where:

$$\begin{aligned} \Delta_1 &= -\alpha^2 + 3\alpha\sigma^2 - \alpha z_0' - z_0'^2, \\ \Delta_2 &= 9\alpha^2\sigma^2 - 3\alpha^2 z_0' - 2\alpha^3 - 9\alpha\sigma^2 z_0' + 27\alpha\sigma^2 x + 3\alpha z_0'^2 + 2z_0'^3, \\ \Delta_3 &= \sqrt[3]{\Delta_2 + \sqrt{4\Delta_1^3 + \Delta_2^2}} \end{aligned}$$

Here, we noticed that except for $z_1'$, there existed only one real root of the equation, denoted as $\hat{z}'_{\mathrm{MAP}}$, which is the MAP estimate of $z'$. Take logarithm of $\hat{z}'_{\mathrm{MAP}}$, we have $\hat{z} = \log(\hat{z}'_{\mathrm{MAP}})$. For normalized scRNA-seq data as shown in equation (5b), the partial derivative with respect to $z$ is:

$$\frac{\partial \mathbb{P}(x|z,\alpha,\psi)\mathbb{P}(z)}{\partial z} \propto \mathrm{e}^{-\frac{(z-z_0)^2}{2\alpha}-\frac{(x-z)^2}{2\sigma^2}}\left(\alpha(x-z)-\sigma^2(z-z_0)\right) \tag{S7}$$

Setting to $0$, we have:

$$\hat{z}_{\mathrm{MAP}} = \frac{sx+\alpha z_0}{s+\alpha} \tag{S8}$$

**The Gillespie algorithm for simulating cell division history in a differentiating population**

| Algorithm 1 - Simulation of cell division history in a differentiating population |
|---|
| 1. Initialize the time $t = t_0$ and cell number of cells of each type $x = x_0$; |
| 2. With the system in state $x$ at time $t$, evaluate all the $a_j(x)$ and their sum $\sum_j a_j(x)$ as follows: $$a_j(x) = x_j R_j$$ where $x_j$ is the number of differentiating or dividing cells in reaction $j$, $R_j$ is the rate of reaction $j$; |
| 3. Effect the next reaction by replacing $t \leftarrow t + \tau$ and $x \leftarrow x + v_j$; |
| 4. Update cell state according to reaction; |
| 5. Record $(x, t)$ as desired; |
| 6. Return to step 1, or end the simulation at population size of 10,000 cells. |

**Table S1 (separate file): The MEGs and their phylogenetic velocity estimations from six phylogeny-resolved scRNA-seq datasets.**

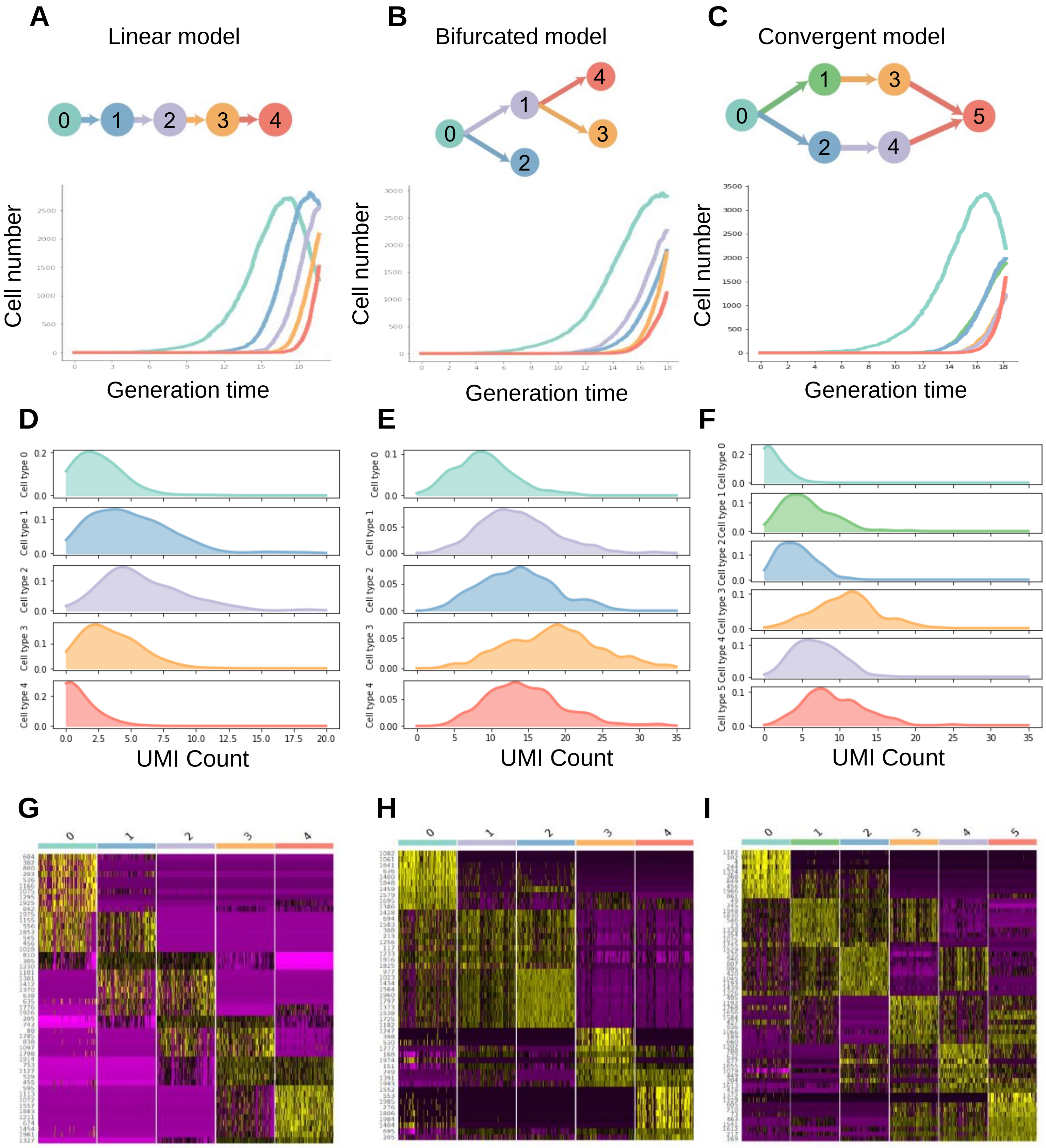


**Fig. S1. Population growth and characteristic gene expressions of different cell types in three differentiation models of simulations**. (**A-C**) Population growth of different cell types in linear differentiation (**A**), bifurcated differentiation (**B**), and convergent differentiation (**C**) models, respectively. (**D-F**) The UMI count distribution of one characteristic gene amongst different cell types in each of the three differentiation models. (**G-I**) Heatmaps showing differential gene expression patterns in the different cell types in three cell differentiation models, respectively.

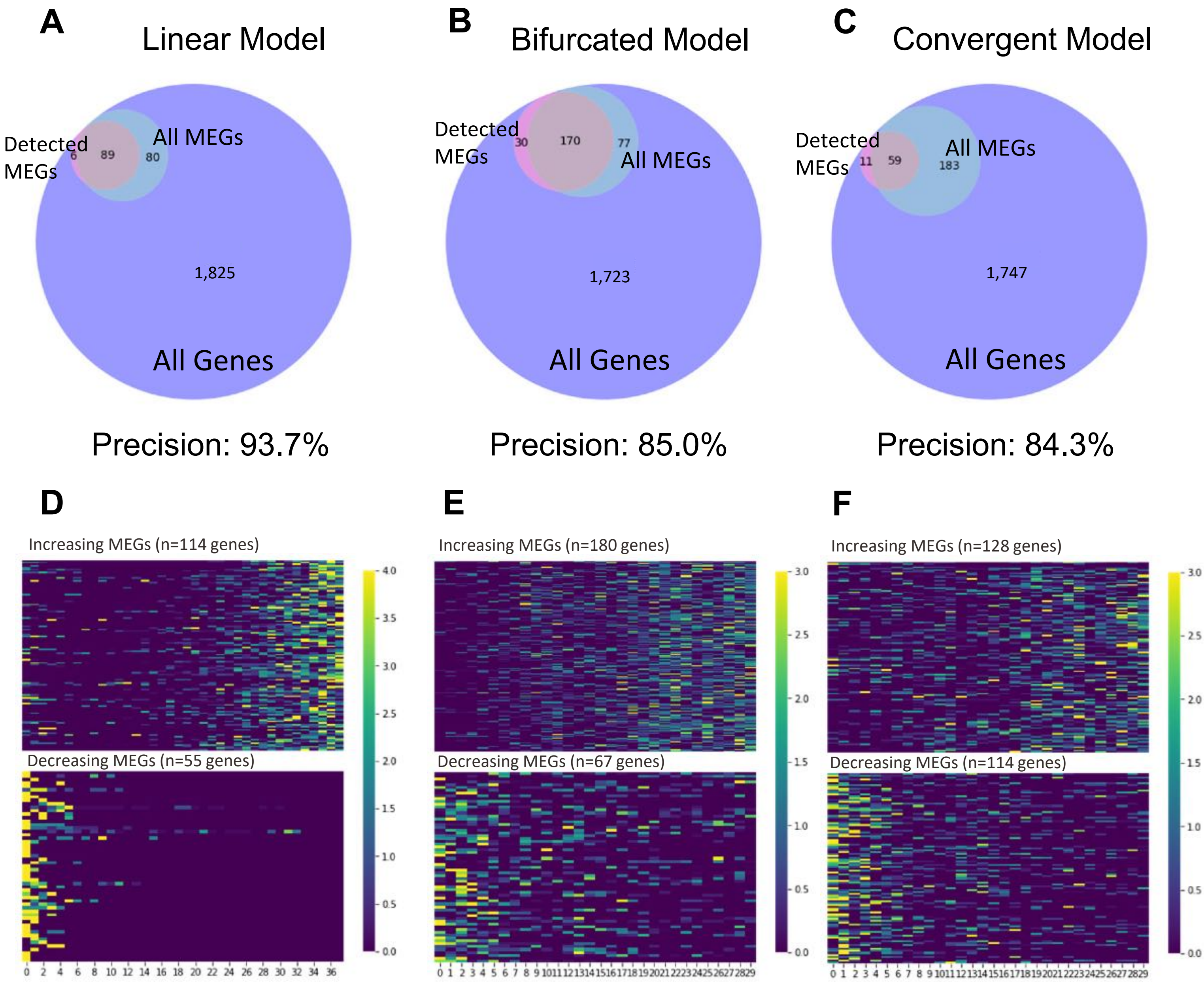


**Fig. S2. High accuracy of our algorithm for identifying MEGs in simulation data. (A-C)** The overlap between the detected MEGs and all real MEGs in the three differentiation models of simulations. (**D-F**) Heatmaps showing the expression dynamics of MEGs along cell divisions in the three differentiation models.

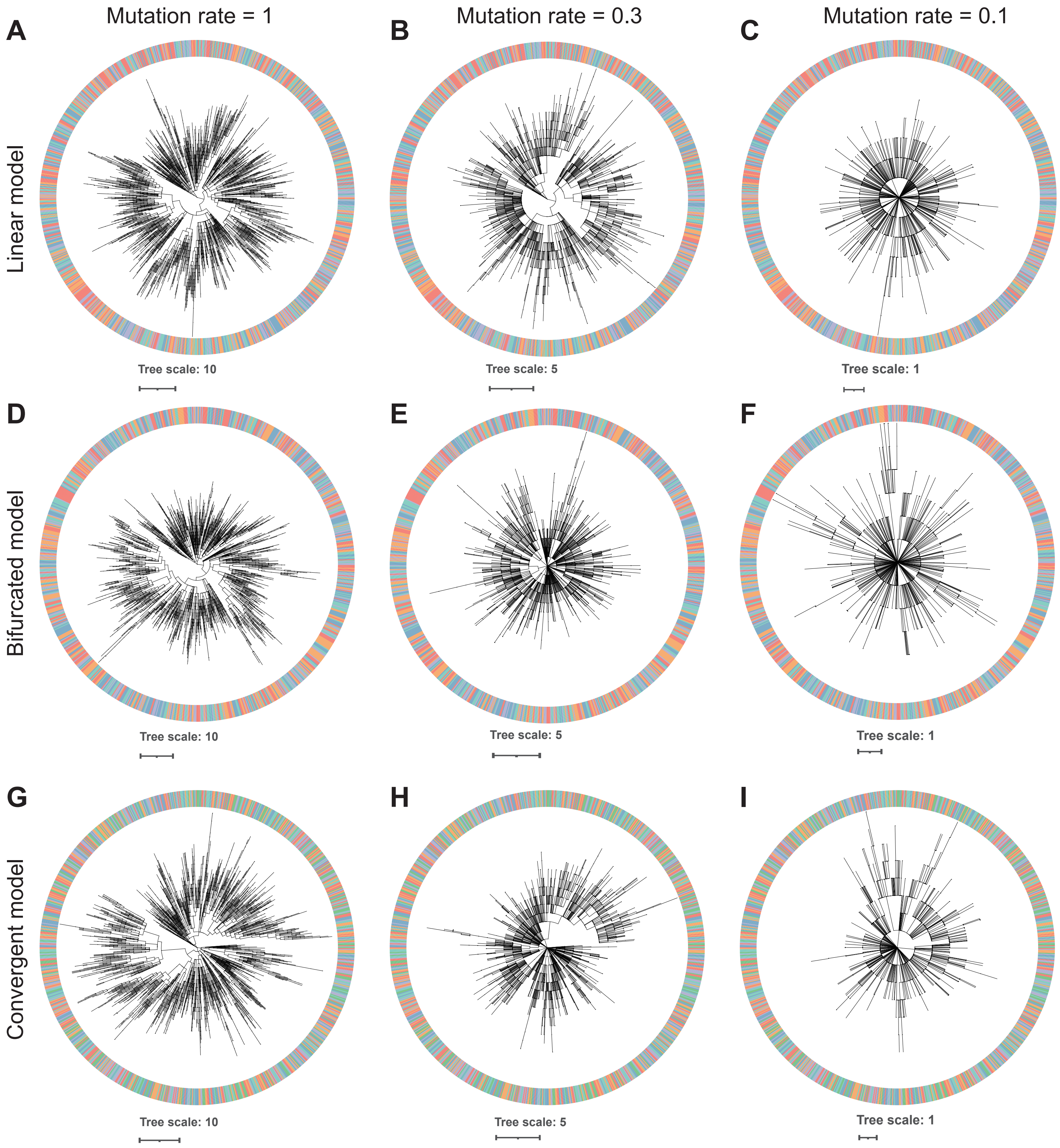


**Fig. S3. Cell phylogenies recorded by simulated mutations.** (**A-C**) Linear differentiation model with mean mutation rate of *u*=1, 0.3, and 0.1, respectively. (**D-F**) Bifurcated differentiation model with mean mutation rate of *u*=1, 0.3, and 0.1, respectively. (**G-I**) Convergent differentiation model with mean mutation rate of *u*=1, 0.3, and 0.1, respectively.

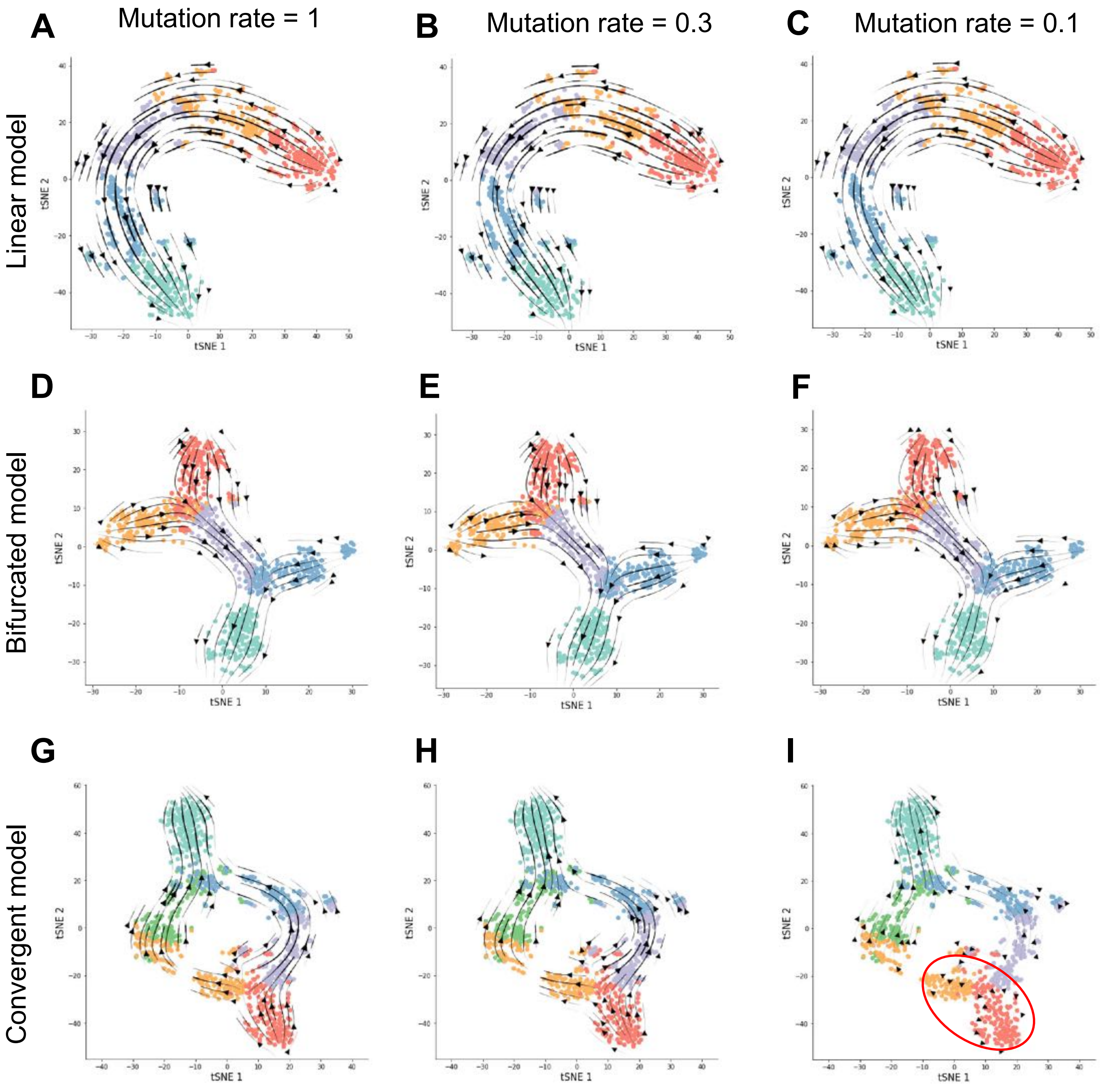


**Fig. S4. PhyloVelo velocity fields in simulation data at different barcoding mutation rates.** (**A-C**) PhyloVelo velocity fields under linear differentiation model with mean mutation rate of *u*=1, 0.3, and 0.1, respectively. (**D-F**) PhyloVelo velocity fields under bifurcated differentiation model with mean mutation rate of *u*=1, 0.3, and 0.1, respectively. (**G-I**) PhyloVelo velocity fields under convergent differentiation model with mean mutRed circle indicates the local cell lineages with poor velocity estimations. ation rate of *u*=1, 0.3, and 0.1, respectively.

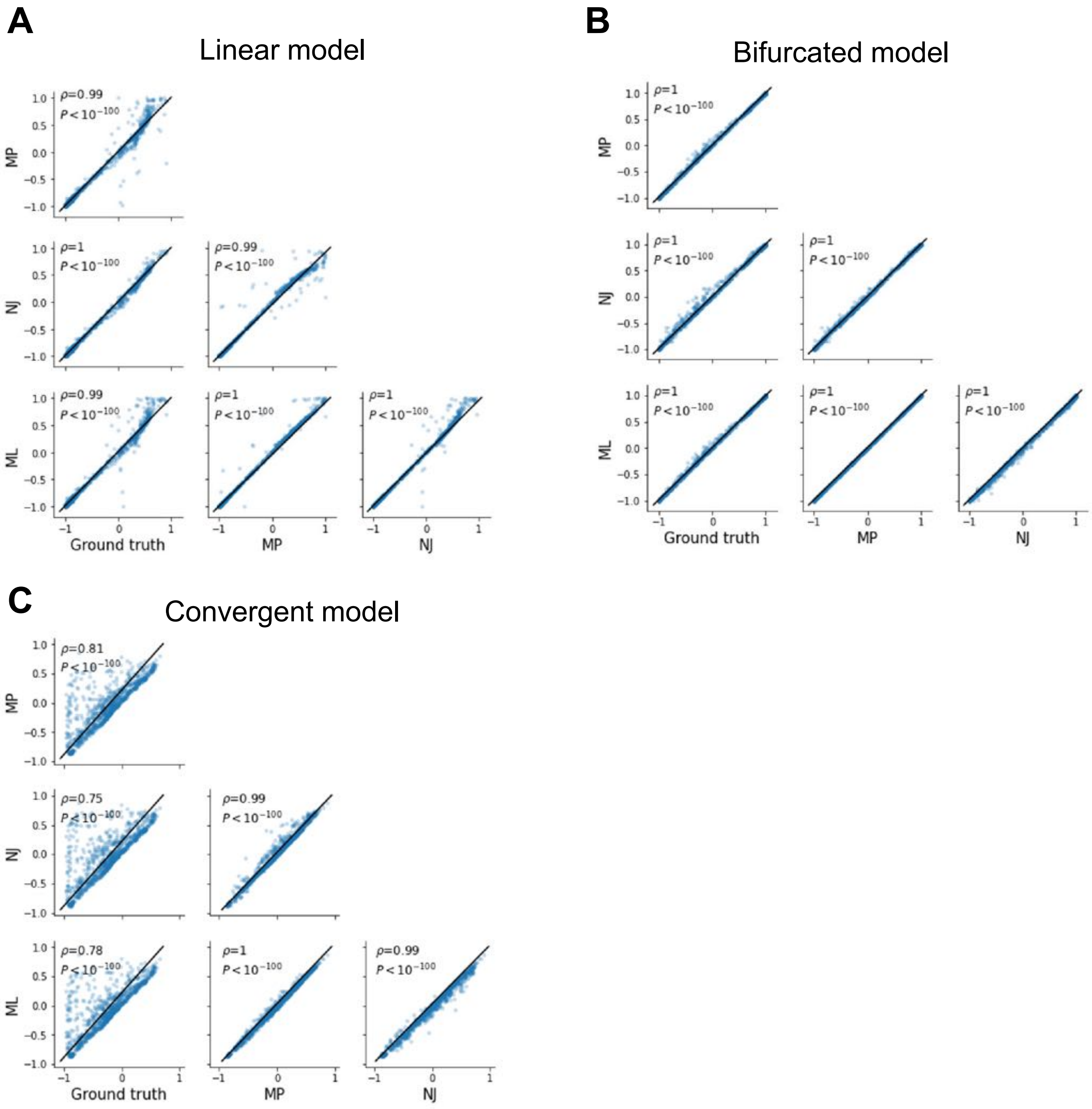


**Fig. S5. PhyloVelo inference is robust to the phylogenetic reconstruction methods.** The pairwise comparison for the directions of phylogenetic velocities based on different phylogenetic reconstruction methods, namely Maximum Likelihood (or ML), Neighbor-Joining (or NJ) or Maximum Parsimony (or MP). (**A**) Linear differentiation. (**B**) Bifurcated differentiation. (**C**) Convergent differentiation. Ground truth refers to the phylogenetic velocity estimated with the ground-truth division history, instead of the mutation-based lineage tree. Here, barcoding mutation rate is $u$=0.3 per cell division.

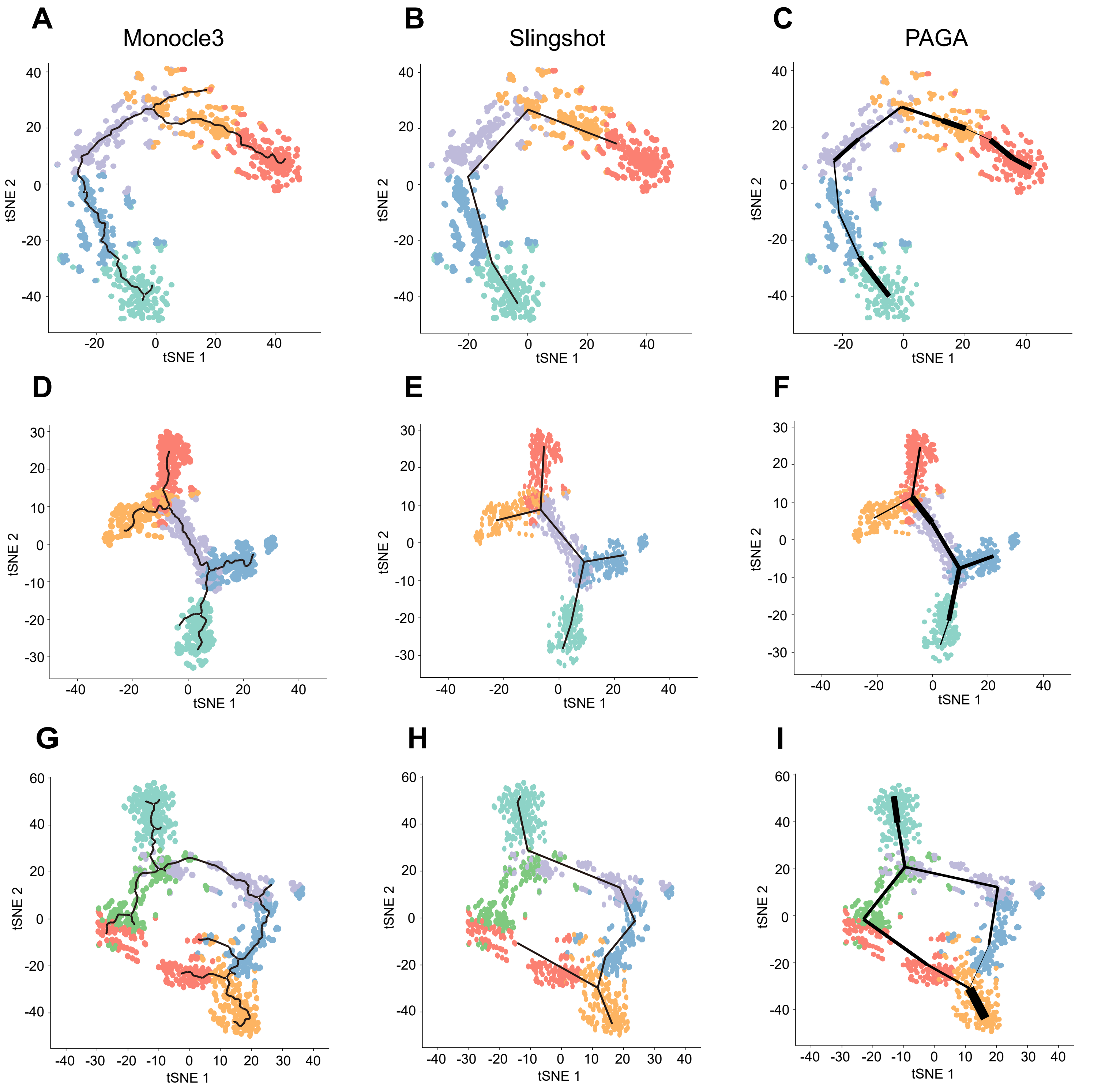


**Fig. S6. Trajectory inference by Monocle3, Slingshot and PAGA, respectively on simulation data.** (**A-C**) Linear differentiation model. (**D-F**) Bifurcated differentiation model. (**G-I**) Convergent differentiation model.

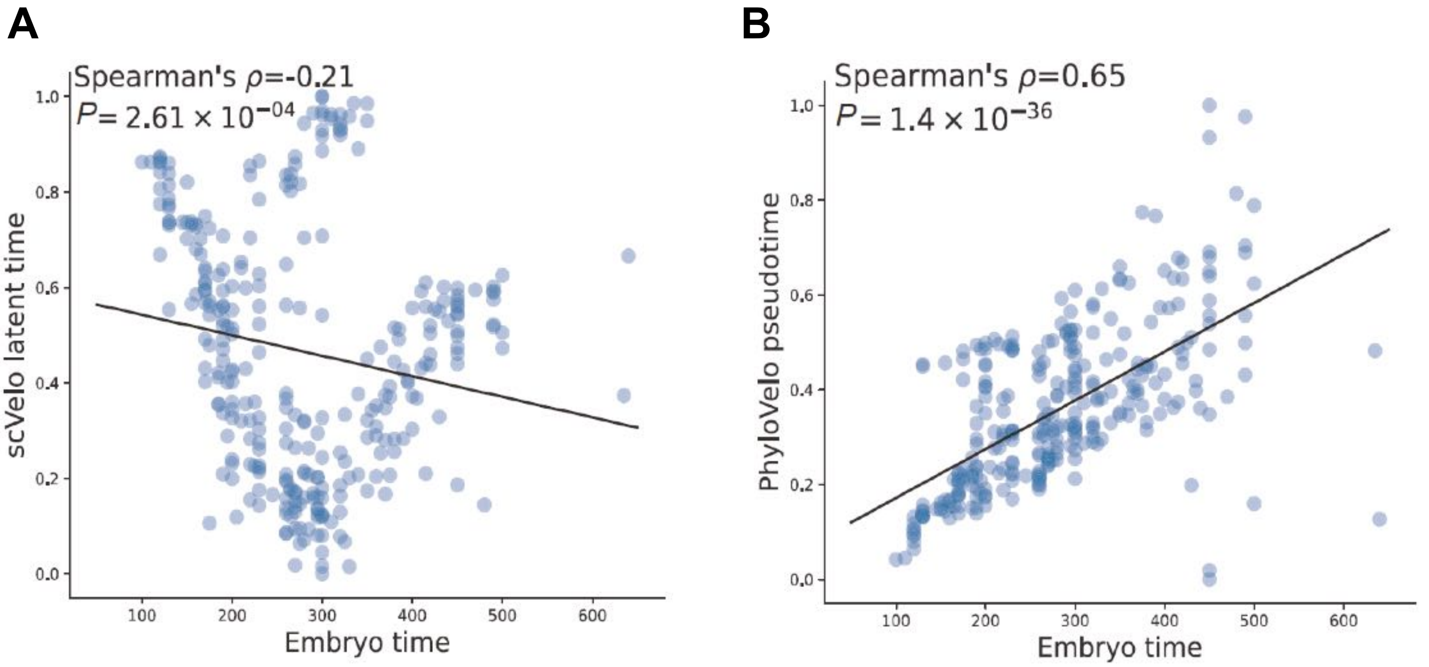


**Fig. S7. The correlation between the velocity latent (or pseudo-) time and *C. elegans* embryo time for the AB lineage cells.** (**A**) scVelo latent time vs embryo time. (**B**) PhyloVelo pseudotime vs embryo time. n=298 cells.

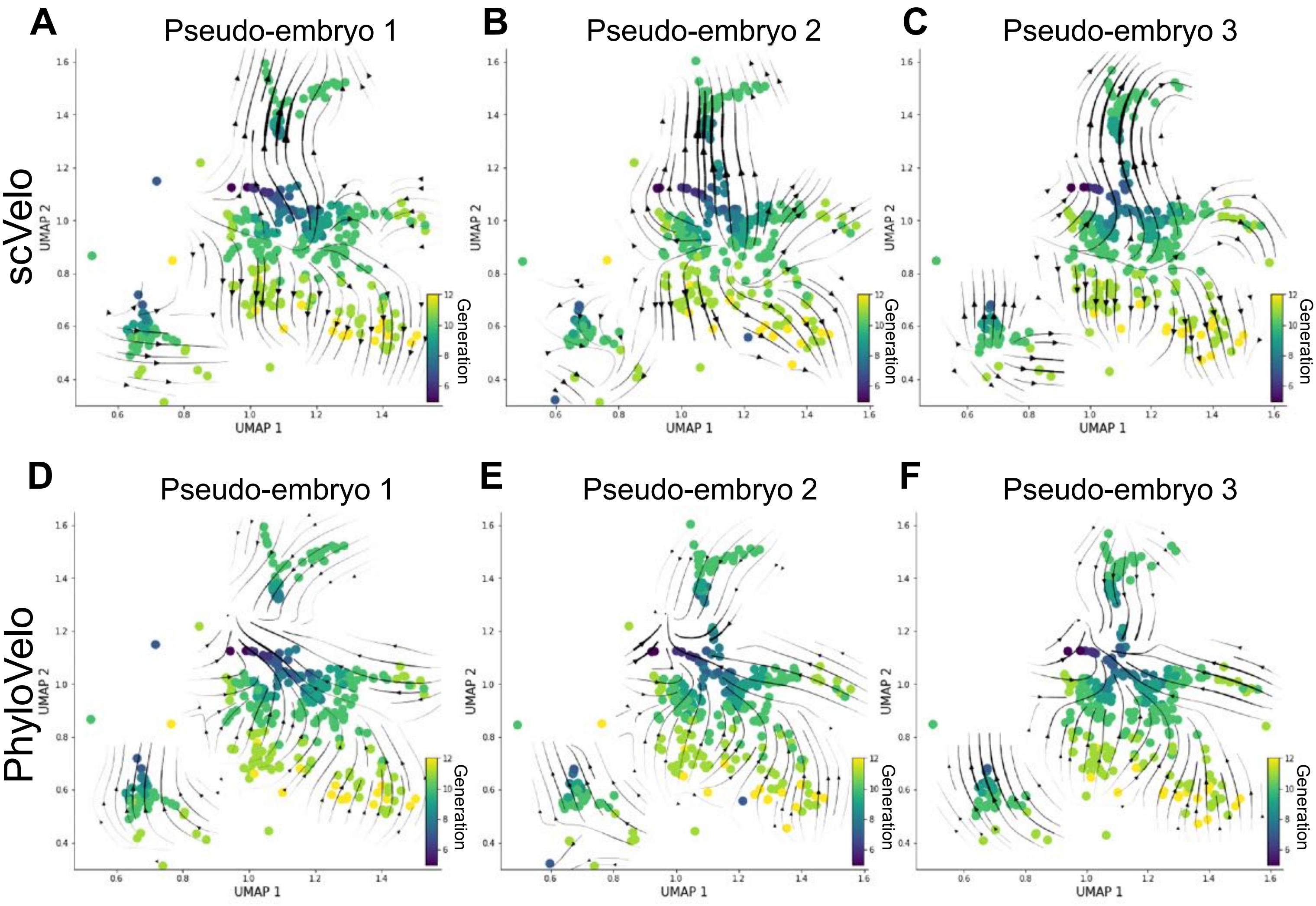


**Fig. S8**. **RNA velocity and PhyloVelo velocity fields in three additional pseudo-embryos of *C. elegans* AB lineage.** (**A-C**) RNA velocity fields by scVelo (dynamic mode). (**D-F**) PhyloVelo velocity fields. A pseudo-embryo refers to the non-repetitive cells (nodes) on the *C. elegans* lineage tree where only one cell from each node of the lineage tree was sampled. The UMAP coordinates are as the original study of Packer *et al*.

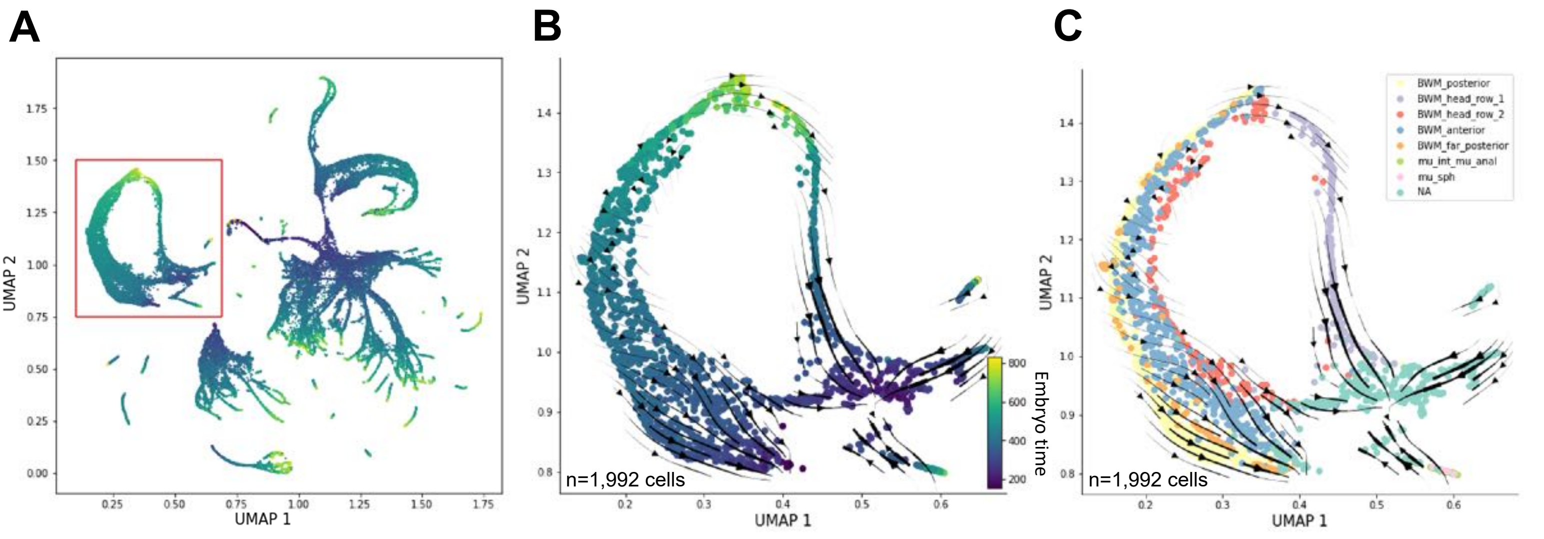


**Fig. S9. PhyloVelo identifies a convergent differentiation in the development of body muscle wall (BMW) in *C. elegans*. (A)** Body muscle wall (BMW) lineage cells were extracted from Packer *et al*., consisting of cells from C, D and MS lineages. (**B-C**) PhyloVelo velocity fields of the pooled BMW lineage cells (n=1,992) from multiple embryos, reconstructed using the MEGs identified from 298 AB lineage cells. Colors are labeled by the embryo time (**B**) or cell types (**C**).

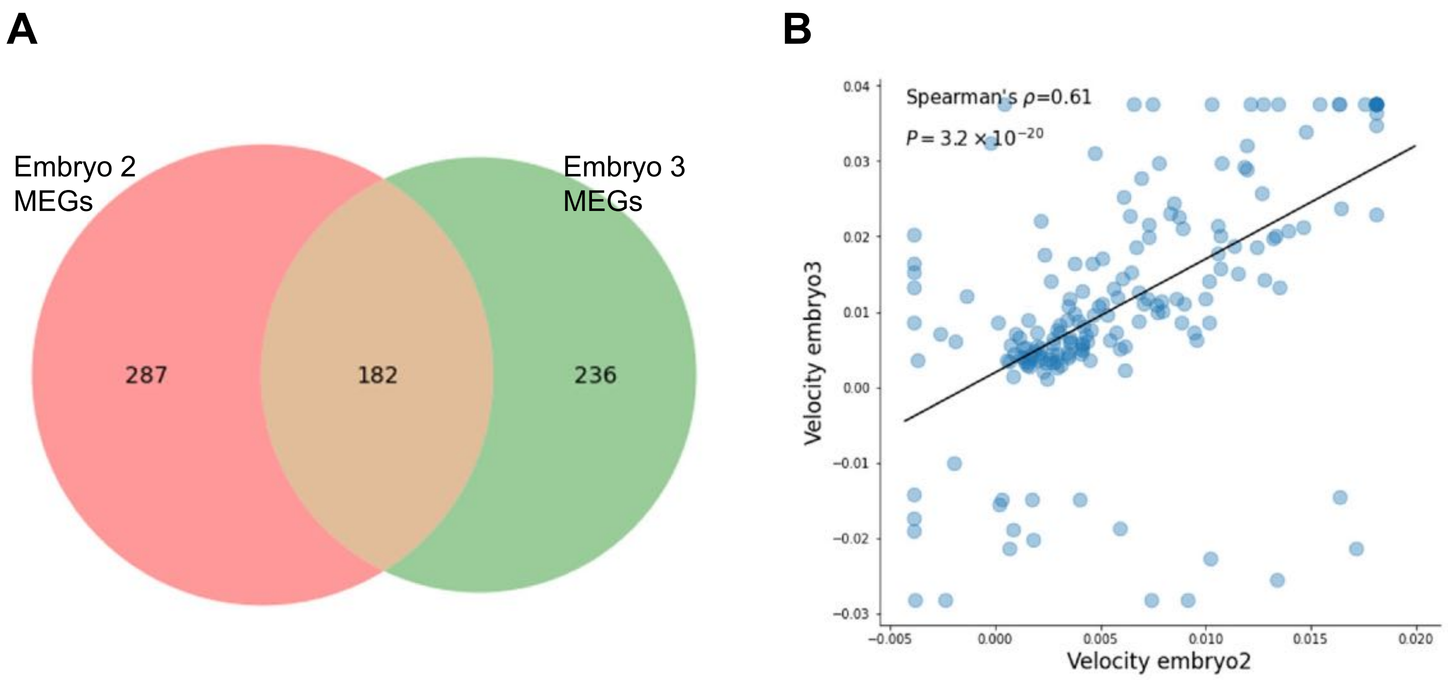


**Fig. S10. High concordance of MEGs identified from two mouse embryos (E8.0/8.5).** (**A**) Venn diagram showing the overlap of MEGs identified from two mouse embryos in the dataset of Chan *et al*., embryo 2 (n=19,017 cells) and 3 (n=10,800 cells). (**B**) The correlation of phylogenetic velocities $\boldsymbol{v}$ for the overlapped MEGs between the two embryos.

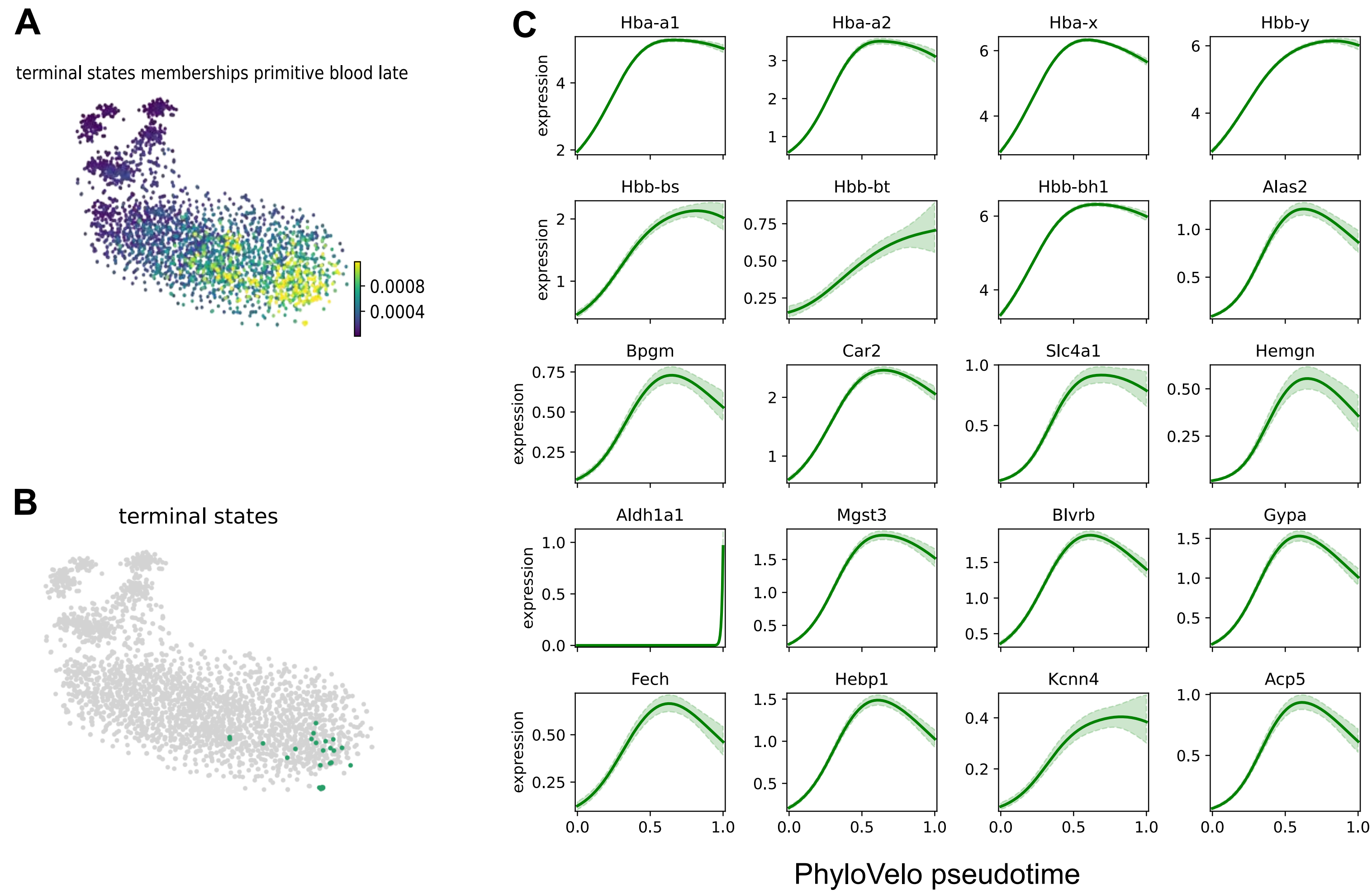


**Fig. S11. Driver genes of erythroid maturation identified by CellRank.** (**A**) The terminal state probability. (**B**) The terminal cell cluster. (**C**) The expression trajectories of driver genes identified by CellRank. PhyloVelo pseudotime was used as the input of CellRank.

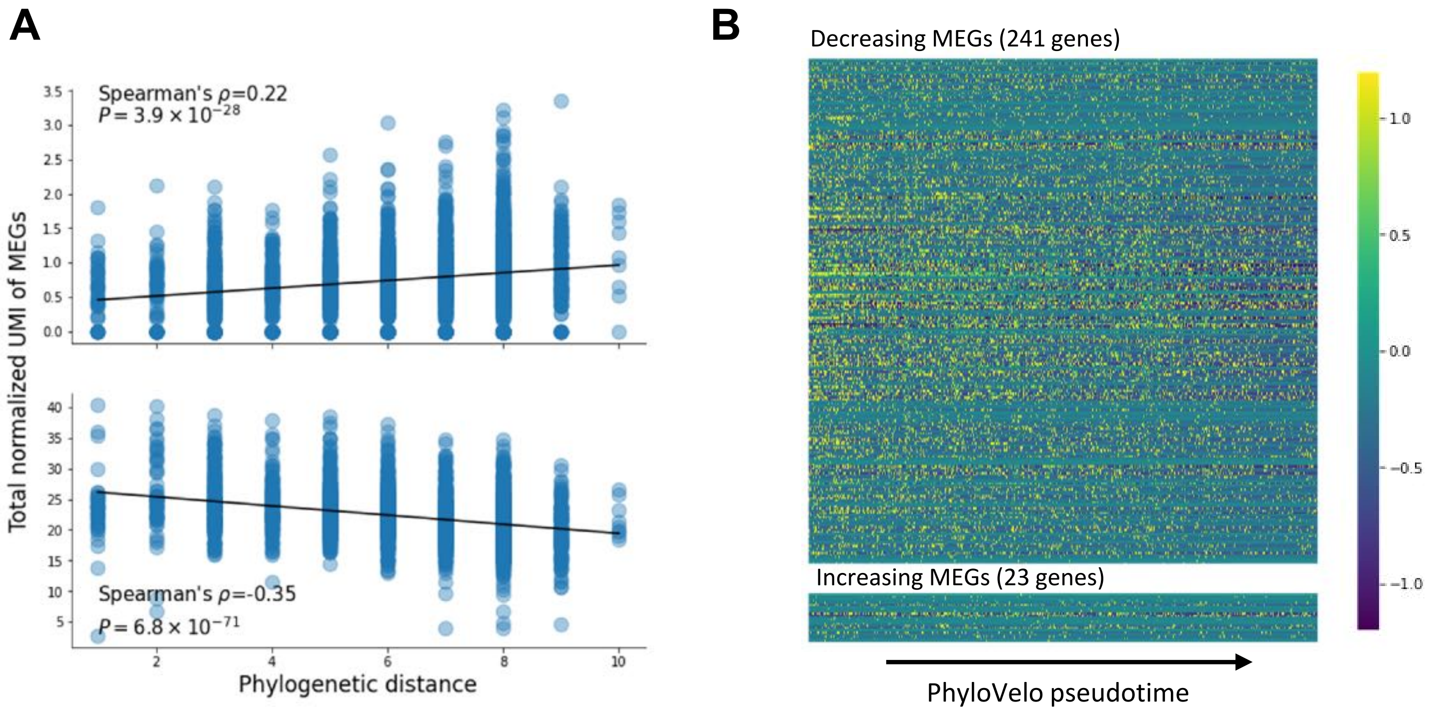


**Fig. S12. The expressions of MEGs identified in the CRISPR lineage tracing dataset of mouse erythroid development. (A)** The total UMI count (normalized) of MEGs changing with the phylogenetic distance from the root. **(B)** Heatmap showing the expression trajectory of MEGs with PhyloVelo pseudotime of erythroid development. Data were from Chan *et al*.

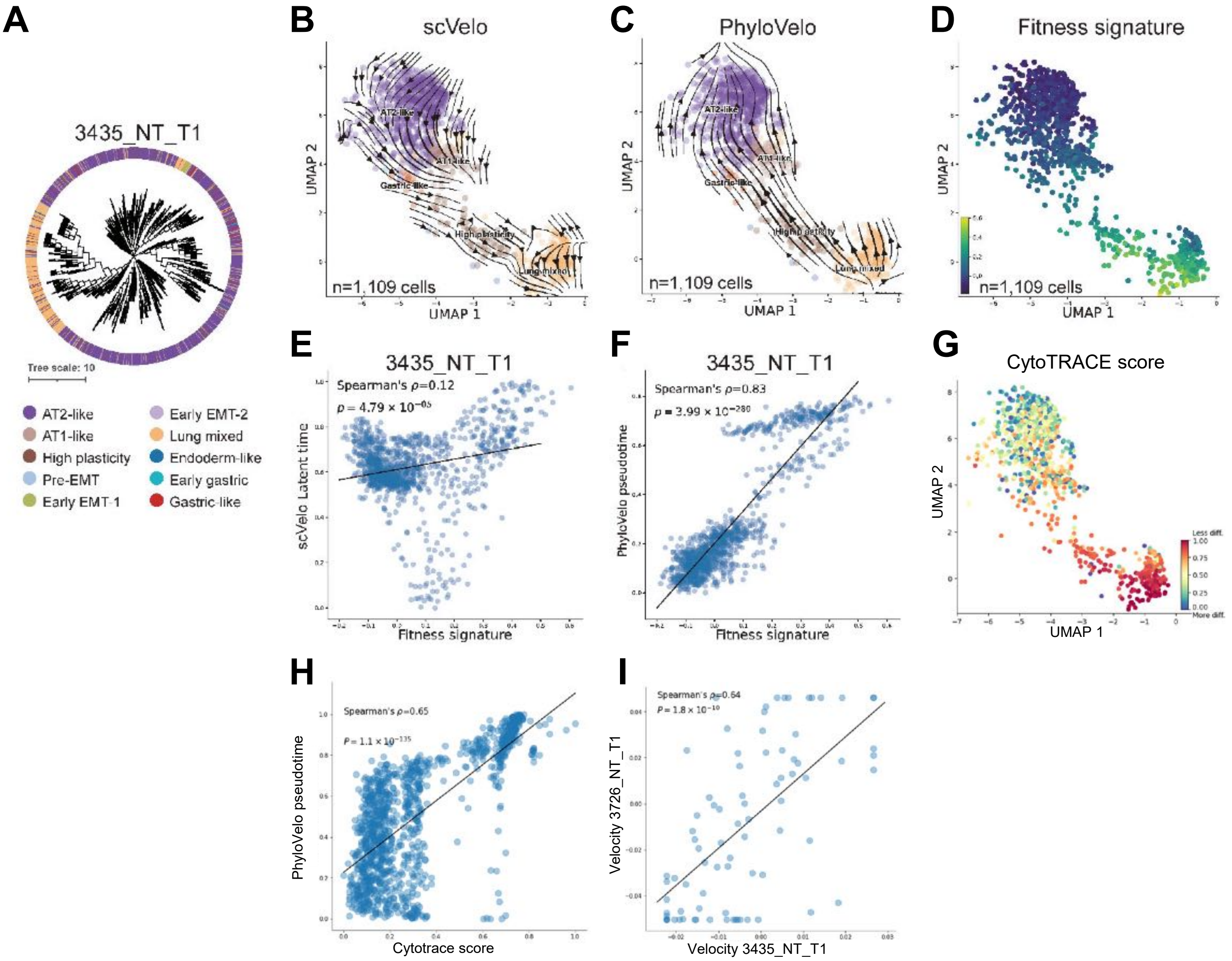


**Fig. S13. PhyloVelo reconstructs the cellular trajectory of lung cancer evolution for 3435_NT_T1.** (**A**) Single-cell phylogenetic tree of primary lung tumor 3435_NT_T1 (n=1,109 cells) from KP ($Kras^{LSL\text{-}G12D/+}$;$Trp53^{fl/fl}$) mouse model. The single-cell RNA data, cell type annotations and lineage tree were obtained from Yang *et al*. (**B**) RNA velocity fields (scVelo - dynamical mode) of tumor 3435_NT_T1. (**C**) PhyloVelo velocity fields of tumor 3435_NT_T1. (**D**) The fitness signatures of single cells for tumor 3435_NT_T1 as defined by Yang *et al*. (**E**) The correlation between scVelo latent time and fitness signatures in tumor 3435_NT_T1. (**F**) The correlation between PhyloVelo pseudotime and fitness signatures in tumor 3435_NT_T1. (**G**) CytoTRACE scores of single cells for 3435_NT_T1. (**H**) The correlation between PhyloVelo pseudotime and CytoTRACE scores. (**I**) The correlation of phylogenetic velocities $\boldsymbol{v}$ for the overlapped MEGs between KP primary tumor 3435_NT_T1 and 3726_NT_T1.

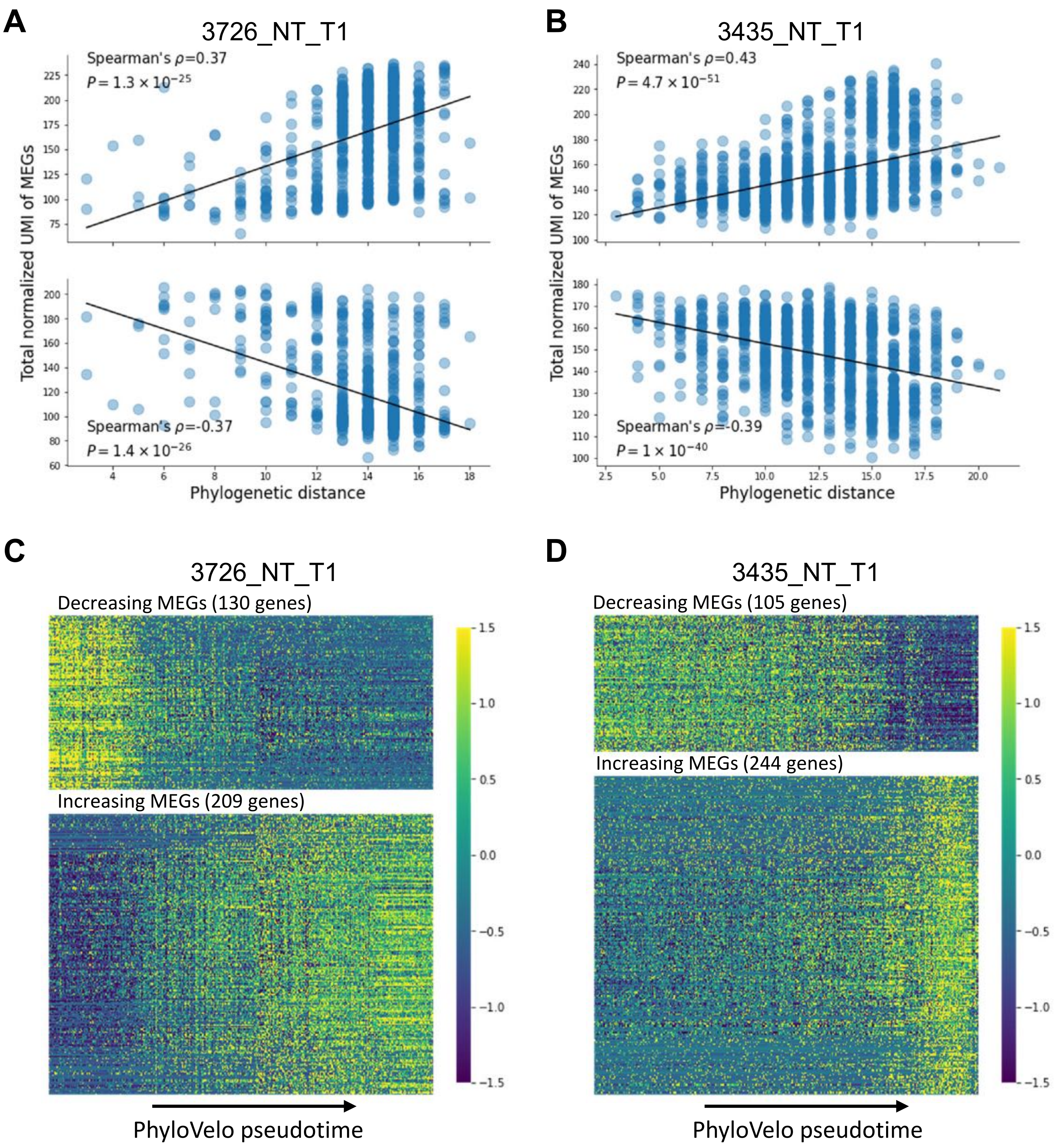


**Fig. S14. Heatmap showing the MEG expression dynamics in KP mouse lung tumors. (A-B**) The total UMI count (normalized) of MEGs changing with the phylogenetic distance from the root in tumor 3726_NT_T1 (**A**) and 3435_NT_T1 (**B**), respectively. **(C-D)** Heatmaps showing the MEGs expression dynamics along PhyloVelo pseudotime in tumor 3726_NT_T1 (**C**) and 3435_NT_T1 (**D**), respectively.

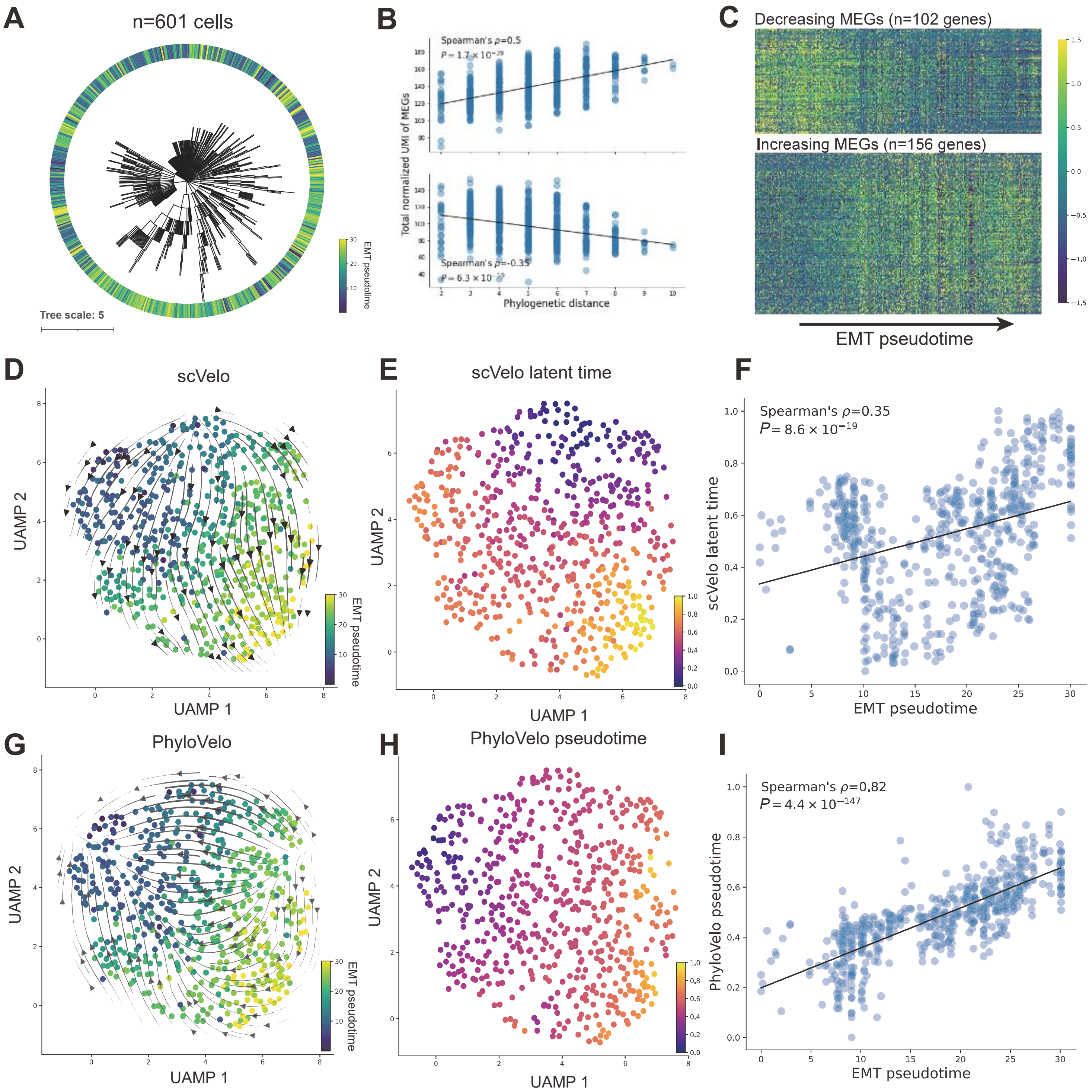


**Fig. S15. The dynamic EMT trajectory in metastatic progression of pancreatic cancer KPCY cells.** (**A**) Phylogenetic tree of 601 non-repetitive terminal cells in tumor subclone M1.1 from Simeonov *et al*. Cell colors are labeled by EMT pseudotime as defined in the original study. (**B**) The total UMI count (normalized) of MEGs changing with the phylogenetic distance from the root. (**C**) Heatmap of MEG expressions (z-score normalized) with EMT pseudotime. (**D**) RNA velocity fields (scVelo - dynamical mode). Cell colors are labeled by EMT pseudotime. (**E**) scVelo latent time. (**F**) The correlation between scVelo latent time and EMT pseudotime. (**G**) PhyloVelo velocity fields. Cell colors are labeled by EMT pseudotime. (**H**) PhyloVelo pseudotime. (**I**) The correlation between PhyloVelo pseudotime and EMT pseudotime.

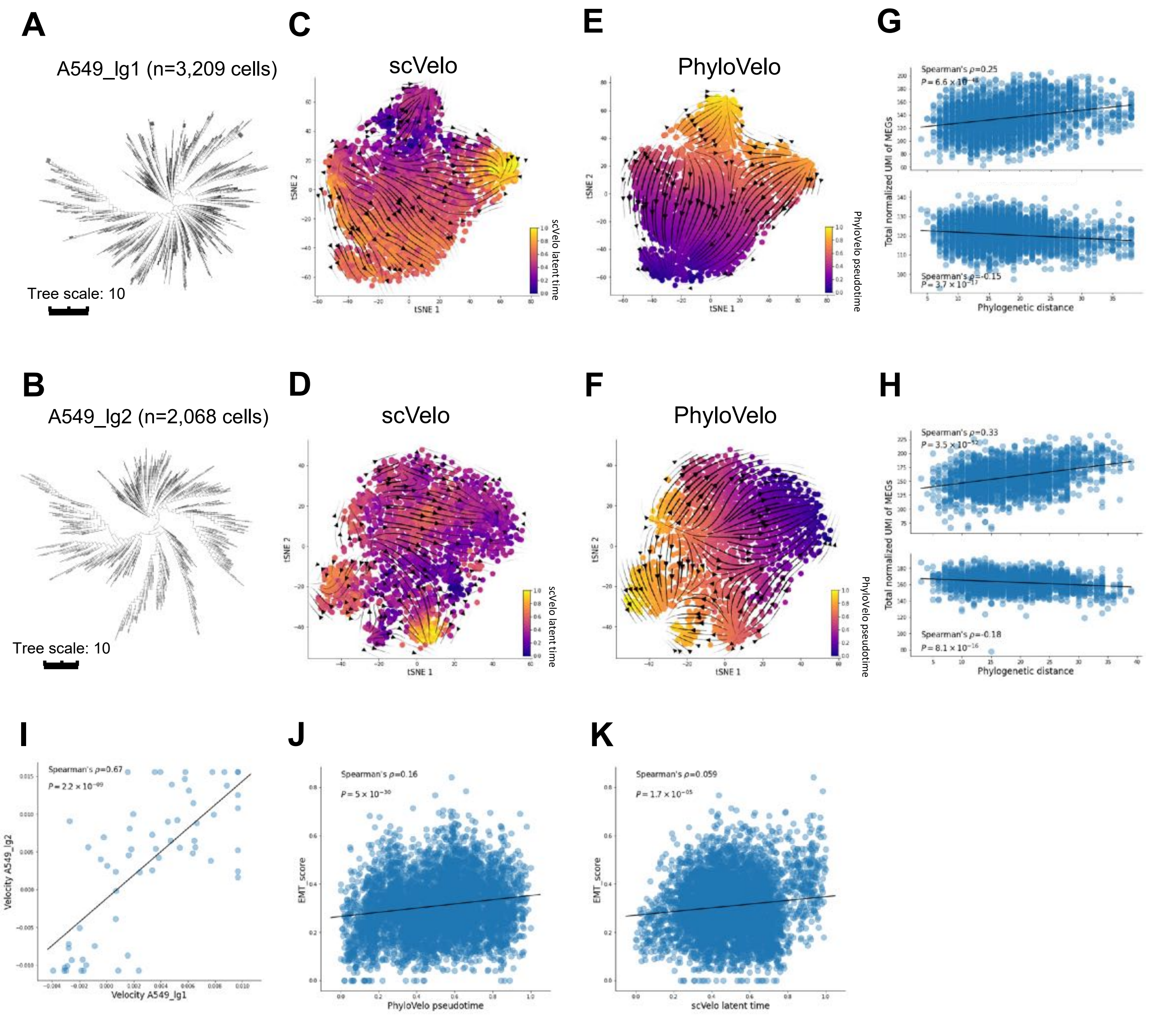


**Fig. S16. Continuous state transitions in the mouse xenograft model of lung cancer cell line A549.** (**A-B**) Phylogenetic trees of the latest clonal cell populations, lg1 and lg2, respectively, obtained from Quinn *et al*. dataset. (**C-D**) RNA velocity fields (scVelo - dynamical mode) for lg1 and lg2, respectively. (**E-F**) PhyloVelo velocity fields for lg1 and lg2, respectively. (**G-H**) The total UMI count (normalized) of MEGs changing with the phylogenetic distance from the root for lg1 and lg2, respectively. (**I**) The correlation of phylogenetic velocities $\boldsymbol{v}$ for the overlapped MEGs between lg1 and lg2. (**J**) The correlation between EMT score and PhyloVelo pseudotime of single cells. (**K**) The correlation between EMT score and scVelo latent time of single cells.

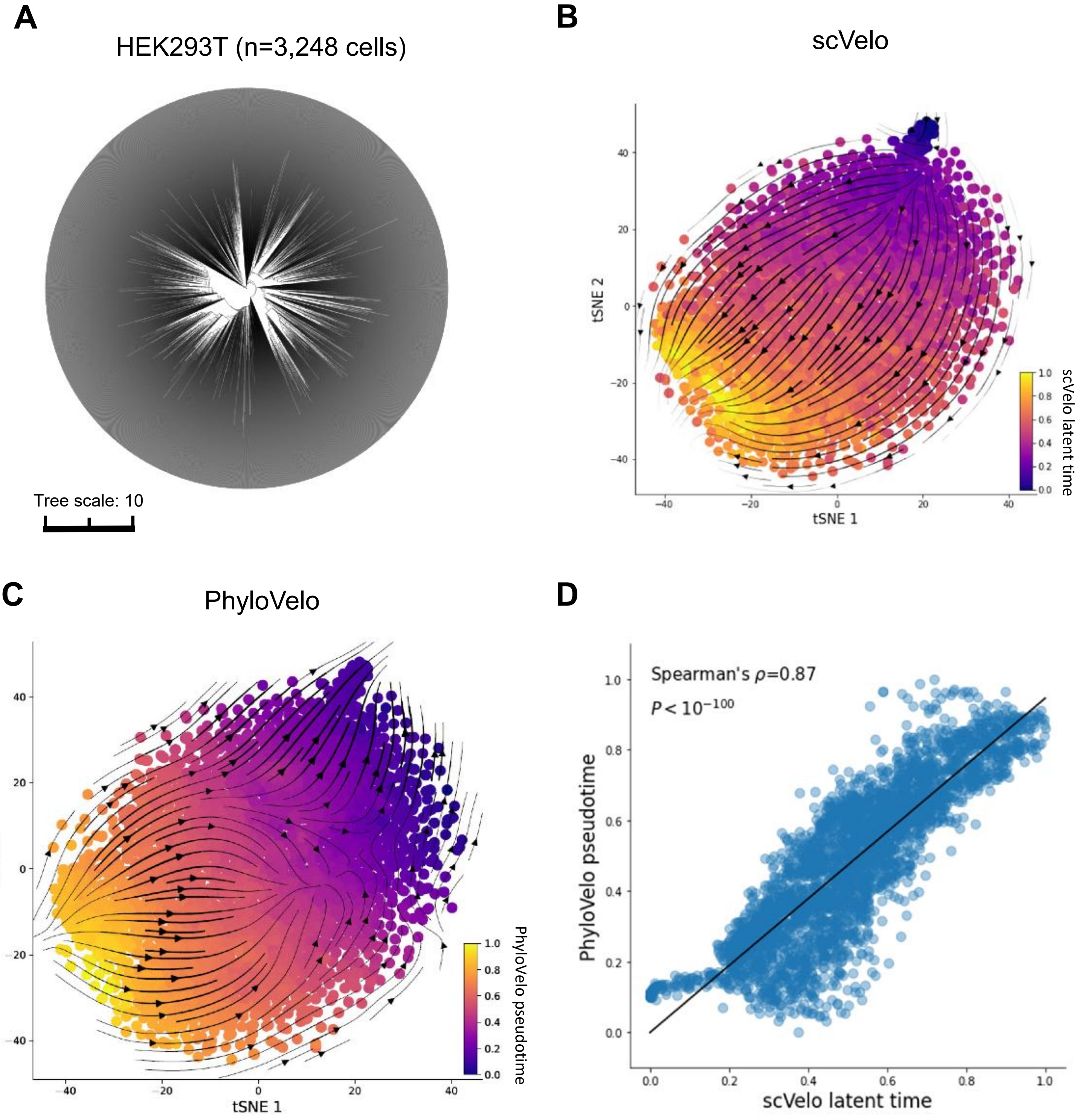


**Fig. S17. Continuous state transitions in the *in vitro* culture of HEK293T cells.** (**A**) The phylogenetic tree of 3,248 HEK293T cells sampled from *in-vitro* culture of a clonal population. (**B**) RNA velocity fields (scVelo - dynamical mode). (**C**) PhyloVelo velocity fields. (**D**) The correlation between PhyloVelo pseudotime and scVelo latent time.

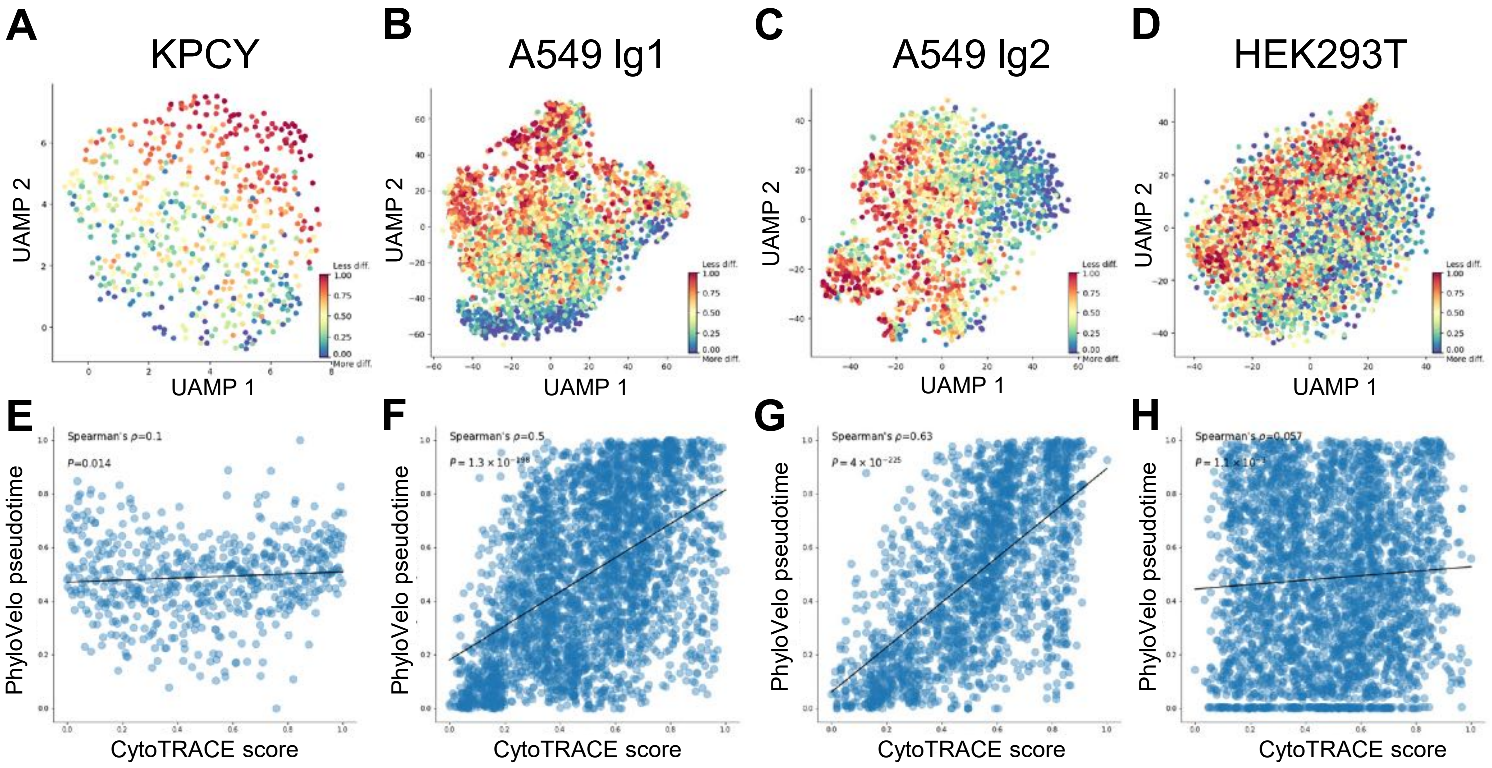


**Fig. S18. Correlation between PhyloVelo pseudotime and CytoTRACE score in four cell line-derived samples.** (**A-D**) CytoTRACE scores in KPCY, A549 lg1, A549 lg2 and HEK293T, respectively. (**E-H**) The correlation between PhyloVelo pseudotime and CytoTRACE score in KPCY, A549 lg1, A549 lg2 and HEK293T, respectively.

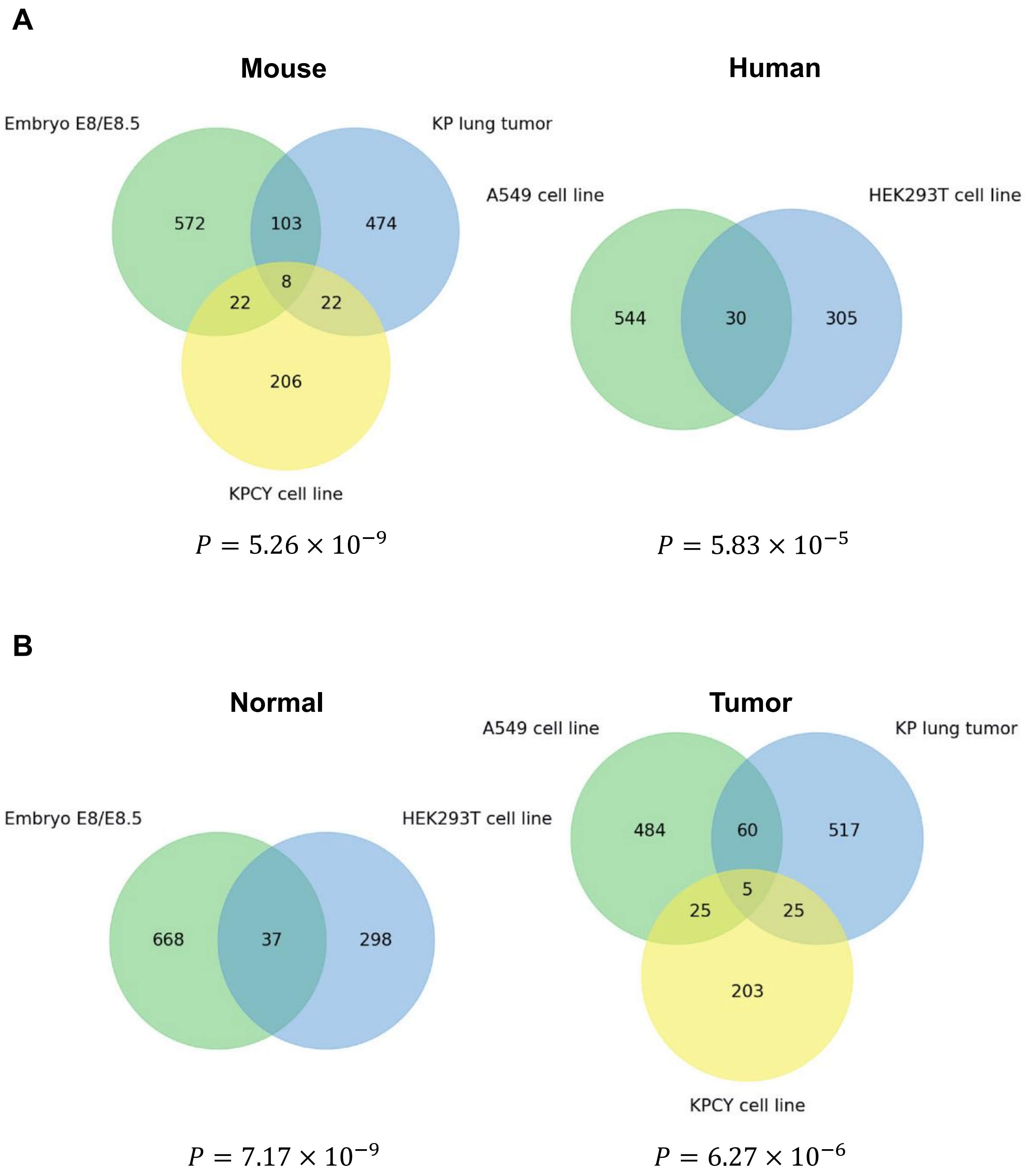


**Fig. S19. Overlap of MEGs across organisms (mouse and human) and tissue or cell types (embryo, primary tumor, cancer and normal cell lines).** (**A**) The overlap of MEGs identified in different datasets as stratified by mouse vs human origin. (**B**) The overlap of MEGs identified in different datasets as stratified by normal vs tumor cells. *P*-values are by SuperExactTest multi-set intersection test.

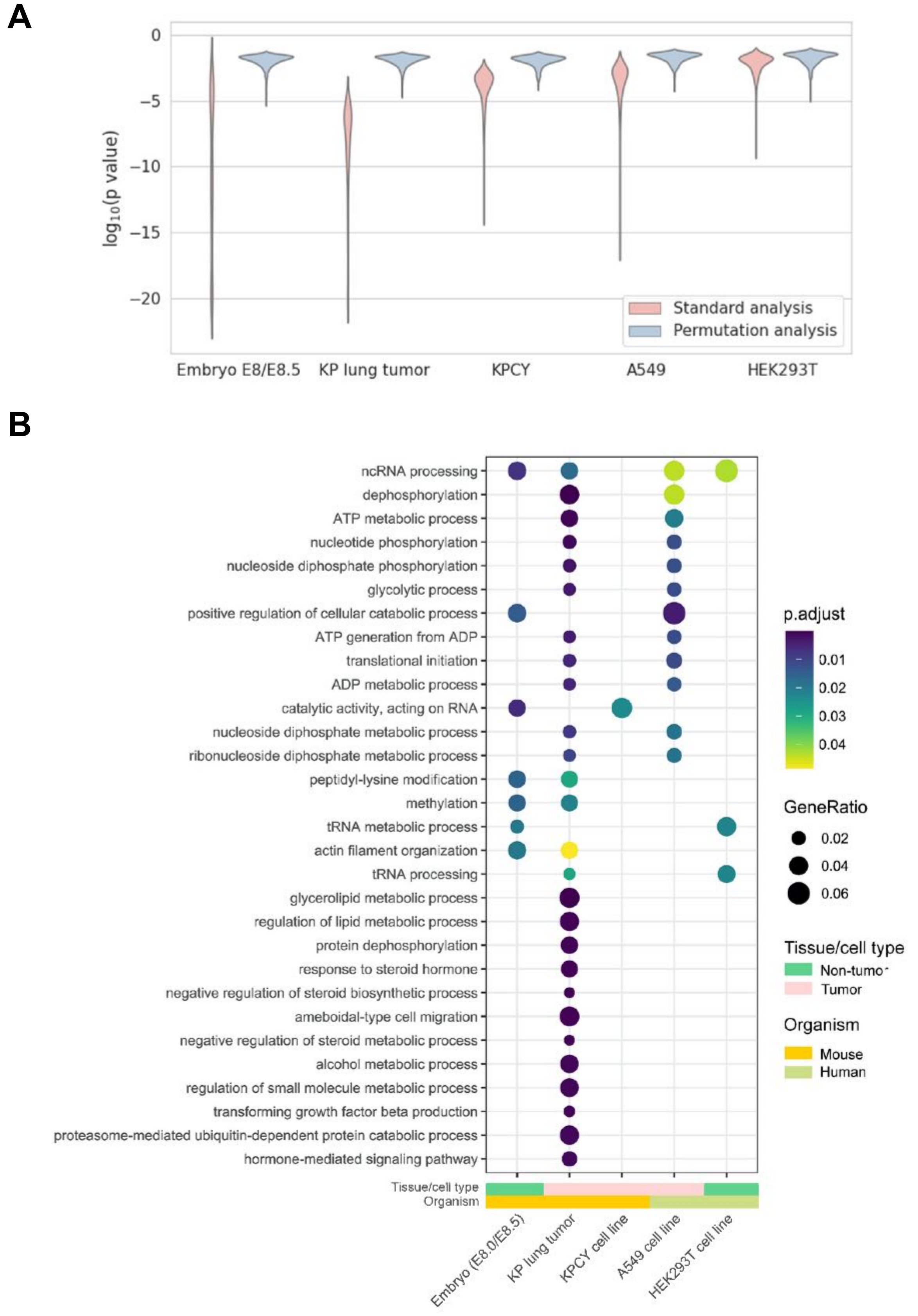


**Fig. S20. Pseudo-MEGs resulting from intrinsic gene expression heterogeneity are functionally different from the genuine MEGs.** (**A**) The *p* values of MEGs in standard and permutation analysis. Permutation analysis was done by randomly shuffling the phylogenetic distances of the cells, followed by the PhyloVelo inference procedure. (**B**) The GO enrichment of pseudo-MEGs across the five CRISPR lineage tracing datasets.
